# PDK1 has a pleiotropic PINOID-independent role in Arabidopsis development

**DOI:** 10.1101/752725

**Authors:** Yao Xiao, Remko Offringa

## Abstract

The 3-Phosphoinositide-Dependent Protein Kinase 1 (PDK1) is a conserved and important master regulator of AGC kinases in eukaryotic organisms. *pdk1* loss-of-function causes a lethal phenotype in animals and yeast. In contrast, only very mild phenotypic defects have been reported for the *pdk1* loss-of-function mutant of the model plant *Arabidopsis thaliana* (Arabidopsis). The Arabidopsis genome contains two *PDK1* genes, hereafter called *PDK1 and PDK2.* Here we show that the previously reported Arabidopsis *pdk1* T-DNA insertion alleles are not true loss-of-function mutants. By using CRISPR/Cas9 technology, we created true loss-of-function *pdk1* alleles, and *pdk1 pdk2* double mutants carrying these alleles showed multiple growth and development defect, including fused cotyledons, a short primary root, dwarf stature, late flowering, and reduced seed production caused by defects in male fertility. Surprisingly, *pdk1 pdk2* mutants did not phenocopy *pid* mutants, and together with the observations that *PDK1* overexpression does not phenocopy the effect of *PID* overexpression, and that *pdk1 pdk2* loss-of-function does not change PID subcellular localization, we conclude that PDK1 is not essential for PID membrane localization or functionality *in planta*. Nonetheless, most *pdk1 pdk2* phenotypes could be correlated with impaired auxin transport. *PDK1* is highly expressed in vascular tissues and YFP:PDK1 is relatively abundant at the basal/rootward side of root stele cells, where it colocalizes with PIN auxin efflux carriers, and the AGC1 kinases PAX and D6PK/D6PKLs. Our genetic and phenotypic analysis suggests that PDK1 is likely to control auxin transport as master regulator of these AGC1 kinases in Arabidopsis.

## Introduction

Protein phosphorylation by protein kinases is a ubiquitous and crucial posttranslational modification in eukaryotic cells. It is involved in almost all cell activities, such as cell division, cell growth and environmental signaling. The AGC kinase family comprises some of the best-characterized protein serine/threonine kinases in eukaryotic cells, such as the founder members cyclic AMP-dependent protein kinase A (PKA) and calcium-dependent protein kinase C (PKC) (Pearce et al., 2010). These kinases play crucial roles in basal cellular functions in lower (yeast) and higher (human/mice) eukaryotes. For example, protein kinase B (PKB/c-Akt) is important in apoptosis inhibition and insulin signaling (Lawlor and Alessi, 2001), whereas p70 ribosomal protein S6 kinase (S6K) plays an important role in mRNA translational control (Pearce et al., 2010; Bahrami-B et al., 2014). AGC kinases themselves are also phosphorylation substrates that can be activated by serine/threonine phosphorylation in the activation loop (T-loop) or in the C-terminal hydrophobic motif of the kinase domain (H-motif) (Chamoto et al., 2010). The 3-Phosphoinositide-Dependent Protein Kinase 1 (PDK1) is a well-established activator responsible for AGC kinase T-loop phosphorylation (Mora et al., 2004; Chamoto et al., 2010).

PDK1 itself is also a conserved member of AGC kinase family, and typically contains a kinase domain with a PDK1-Interacting Fragment (PIF)-binding pocket at its N-terminus and a PH domain at the C-terminus (Biondi et al., 2000; Frödin et al., 2002). Other AGC kinases have a C-terminal hydrophobic PIF motif, and the interaction with the PIF binding pocket in PDK1 enhances their activation by phosphorylation. The PH domain is essential for PDK1 plasma membrane recruitment and kinase activity in mammals. Binding of the PH domain to the phospholipid phosphatidylinositol (3,4,5)-trisphosphate [PtdIns(3,4,5)P3] at the plasma membrane triggers PDK1 dimer to monomer conversion and phospho-activation (Alessi et al., 1997; Ziemba et al., 2013). PDK1 was originally named PtdIns(3,4,5)P3-dependent protein kinase 1 (Alessi et al., 1997), but the name was changed when PtdIns(3,4)P2, PtdIns3P and PtdIns(4,5)P2 also appeared to bind its PH domain (Currie et al., 1999; Deak et al., 1999). *Arabidopsis thaliana* PDK1 (AtPDK1) binds to an even broader selection of phospholipids *in vitro* (Deak et al., 1999). Nevertheless, the two most important phospholipids for mammalian PDK1, PtdIns(3,4,5)P3 and PtdIns(3,4)P2, have not been identified in *Arabidopsis thaliana* (Arabidopsis) (Heilmann, 2016), and AtPDK1 activity has been reported to be controlled by PtdIns(4,5)P2 and phosphatidic adic (PA) (Anthony et al., 2004). Arabidopsis has two highly homologous *PDK1* genes, At5g04510 (*AtPDK1.1*) and At3g10540 (At*PDK1.2*), and for convenience reasons we renamed them to respectively *PDK1* and *PDK2*.

Interestingly, the two yeast PDK1 orthologs Pkh1 and −2, which lack a PH domain, still have the ability to phosphorylate AGC kinases (Casamayor et al., 1999; Niederberger and Schweingruber, 1999; Voordeckers et al., 2011). *Physcomitrella patens* PDK1 (PpPDK1), which also lacks a PH domain, is able to rescue the lethal phenotype of the yeast *pkh1 pkh2* double mutant. This indicates that the PH domain is not required in all eukaryotes or full PDK1 functionality (Dittrich and Devarenne, 2012a). Besides for yeast, complete loss-of-function of *PDK1* is also lethal for fruit flies and mice (Lawlor et al., 2002; Rintelen et al., 2002). In plants, several methods have been employed in different species to analyze PDK1 function *in planta*. Virus-induced gene silencing (VIGS) has been used to knock down tomato *PDK1*, whereas *Tos17* transposon mutagenesis or homologous recombination has been used in rice or *Physcomitrella patens*, respectively. However, in tomato the claimed cell death phenotype made *PDK1* knock-out expression unprovable (Devarenne et al., 2006), and in rice the *Tos17* insertion only led to a knock down of *PDK1* expression (Matsui et al., 2010; Dittrich and Devarenne, 2012a). Deletion of *PDK1* in *Physcomitrella patens* was not lethal, but *pdk1* knock-out mutants showed strong developmental defects (Dittrich and Devarenne, 2012a). In Arabidopsis, three combinations of *pdk1 pdk2* T-DNA insertion alleles have been reported to show altered sensitivity to *Piriformospora indica* induced growth promotion, and a weak developmental defect resulting in reduced silique length (Camehl et al., 2011; Scholz et al., 2019). Inhibition of *PDK1* expression in Arabidopsis cell suspensions using RNAi technology delivered no mutant cell phenotype (Anthony et al., 2004).

In contrast to the lack of a clear *in planta* role for PDK1, all Arabidopsis AGC kinases phosphorylated by PDK1 *in vitro*, including PINOID (PID), Oxidative Signal-Inducible1(OXI1), UNICORN (UCN) and most AGC1 family members, do play key roles in plant development and defense (Anthony et al., 2004, 2006; Zegzouti et al., 2006a, 2006b; Devarenne et al., 2006; Camehl et al., 2011; Enugutti et al., 2012; Gray et al., 2013; Scholz et al., 2019). PID phosphorylates PIN auxin efflux carries to control their polarity and thereby direct the auxin flux (Christensen et al., 2000; Benjamins et al., 2001; Friml et al., 2004; Kleine-Vehn et al., 2009; Dhonukshe et al., 2010; Huang et al., 2010). OXI1 plays a dual role in regulating both root hair growth and the basal immune response against virulent pathogen infection (Anthony et al., 2004; Rentel et al., 2004; Anthony et al., 2006; Petersen et al., 2009; Matsui et al., 2010; Camehl et al., 2011). UCN was recently shown to be a phosphorylation target of PDK1 *in vitro*, but genetic evidence suggests that UCN negatively regulates PDK1 at the post-transcriptional level to control planar growth (Scholz et al., 2019). The other established PDK1 targets all belong to the AGC1 protein kinases family (Galván-Ampudia and Offringa, 2007; Rademacher and Offringa, 2012), which has been well-characterized during the past decade. The D6 protein kinases (D6PKs, including D6PK/AGC1.1, D6PKL1/AGC1.2, D6PKL2/PK5, D6PKL3/PK7), PROTEIN KINASE ASSOCIATED WITH BRX (PAX/AGC1.3), PAX LIKE (PAXL/AGC1.4) and AGC1-12 all have been shown to phosphorylate PIN proteins to enhance auxin transport activity (Zourelidou et al., 2009; Willige et al., 2013; Barbosa et al., 2014, 2018; Haga et al., 2018; Marhava et al., 2018). The tomato ortholog of PAX, also known as AvrPto-dependent Pto-interacting protein 3 (Adi3), negatively controls plant cell death caused by pathogen attack (Devarenne et al., 2006; Gray et al., 2013). AGC1.5 and AGC1.7 control polar growth of pollen tubes by phosphorylating RopGEFs. (Zhang et al., 2009; Li et al., 2018), and ROOT HAIR SPECIFIC 3 (RSH3/AGC1.6) specifically regulates root hair morphology (Won et al., 2009). The disproportion between the *in vivo* data on the functions of the different Arabidopsis AGC kinases that are established *in vitro* phosphorylation targets of AtPDK1, and the small role that AtPDK1 itself seems to play in development based on the *pdk1 pdk2* double mutant phenotype, made us reinvestigate the published data on *AtPDK1*.

PID has been reported as one of the prime targets of PDK1 (Zegzouti et al., 2006a), but the *pdk1 pdk2* double mutant lacks the typical *pid* loss-of-function phenotypes. We therefore generated Arabidopsis lines overexpressing *PDK1* (*PDK1*ox), and found that seedlings of these lines lacked the strong phenotypes observed in seedlings overexpressing *PID (PIDox*). These results suggest that either PDK1 requires activation and that this is not triggered in the *PDK1ox* seedlings, or that it is not rate limiting for PID activity. Next, we re-analyzed the published *pdk1* and *pdk2* T-DNA insertion alleles. Based on RT-PCR experiments, the two *pdk2* alleles appeared to represent true loss-of-function mutants. However, functional *PDK1* mRNA was still detectable in the three *pdk1* alleles, explaining the lack of strong phenotypes in the *pdk1 pdk2* double mutant combinations. Using CRISPR/Cas9, we generated several true *pdk1* loss-of-function mutant alleles, which when combined with the *pdk2* T-DNA insertion allele did display strong growth and developmental defects. The mutant phenotypes indicate a pleiotropic, but PID-independent role for PDK1 in plant development as regulator of auxin transport.

## Results

### *PDK1ox* and *PIDox* seedlings do not share phenotypes

The key defects caused by *PIDox* in Arabidopsis are agravitropic seedling growth and collapse of the main root meristems as a result of redirected polarity of PIN-mediated auxin transport (Figure 1G,H, J) (Benjamins et al., 2001; Friml et al., 2004). In view of the model that *PDK1* regulates *PID* kinase activity (Zegzouti et al., 2006a), we expected *PDK1ox* to cause similar phenotypes as *PIDox*. More than thirty independent *p35S::YFP:PDK1* or *p35S::PDK1* transgenic lines were selected and T2 seedlings grown on vertical agar plates showed normal gravitropic growth. Five single locus homozygous T3 lines with different *PDK1* overexpression levels were subsequently selected for further phenotype observation and quantification (Figure 1I). All of the representative *PDK1ox* lines showed normal gravitropic seedling growth and no collapse of the main root meristem was observed (Figure 1). Roots of *p35S::YFP:PDK1#5.4* and *p35S::YFP:PDK1#9.6* seedlings were even slightly longer than wild-type roots (Figure 1H), however, this phenotype did not clearly correlate with the *PDK1* overexpression level (Figure 1I). Also mature *PDK1ox* plants developed and flowered like wild-type Arabidopsis plants.

**Figure 1.**
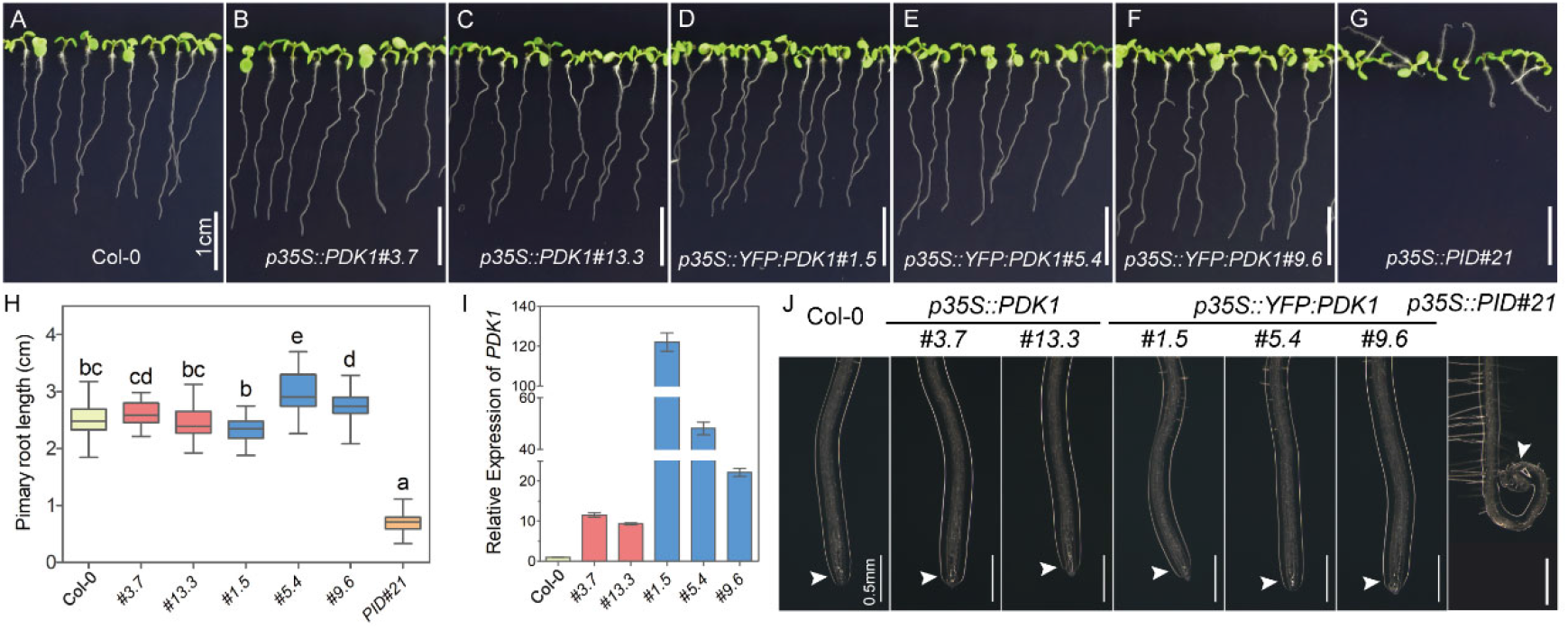
Seedling and root phenotype of *PDK1* and *PID* overexpression lines. A-G, Representative 7-day-old seedlings for indicated lines. Please note that only *p35S::PID*#21 seedlings show agravitropic growth. Scale bars represent 1cm. H, Box plot with Min/Max whiskers showing the quantification of the primary root length of 7-day-old seedlings of Arabidopsis wild type (Col-0), *p35S::PDK1* lines #3.7 and 13.3 (red box), *p35S::YFP:PDK1* lines #1.5, 5.4 and 9.6 (blue box), and *p35S::PID* line #21 The results are from a single experiment (n>36 per line), but similar results were obtained in 3 experimental repeats. Lower case letters indicate statistically different groups (p < 0.05), as determined by a one-way ANOVA followed by Tukey’s test. I, PDK1 expression levels in Col-0 and in the *p35S::PDK1* and *p35S::YFP:PDK1* lines used in H. The bar graph shows the mean value ± SEM. J, Representative images showing a detail of the root tip phenotype of seedlings in A-H. White arrow heads point out collapsed (*p35S::PID*#21) or normal root meristems (all other lines). Scale bar indicates 0.5 mm.

The above results suggest that PDK1 is not rate limiting for endogenous PID activity. However, we cannot exclude that the PDK1 kinase itself requires signaling to be activated, and that therefore its overexpression does not lead to additional phenotypes under normal growth conditions.

### CRISPR/Cas9-generated mutant alleles indicate a central role for PDK1 in development

To obtain further indications for the proposed role of PDK1 as upstream regulator of PID, we re-assessed the previously described *pdk* loss-of-function mutant alleles. Three *pdk1* and two *pdk2* T-DNA insertion alleles have been reported to be loss-of-function mutants (Camehl et al., 2011; Scholz et al., 2019). Neither *pdk1* nor *pdk2* single mutants showed any noticeable phenotype, and different double mutant combinations of the *pdk1* and *pdk2* alleles only showed a mild reduction in silique length and plant height (Camehl et al., 2011; Scholz et al., 2019).

Two *pdk2* alleles, *pdk2-1* and *pdk2-4*, were confirmed to be true knock-out mutants by RT-PCR analysis (Figure 2A, B). However, in contrast to published data, the *pdk1-c* allele appeared to produce a full length mRNA (Figure 2A, B) (Camehl et al., 2011; Scholz et al., 2019), whereas the *pdk1-a* and *pdk1-b* alleles produced a partial or mutated mRNA (Figure 2A, B, Figure S1), leading to the production of a PDK1 kinase lacking its PH domain (Figure S1). Previous studies have suggested that a PH domain may not be essential for PDK1 function in plants (Dittrich and Devarenne, 2012b). When we tested the kinase activity of PDK1 lacking a PH domain *in vitro*, it showed very high autophosphorylation activity (Figure 2C). Based on these findings, we concluded that the three published *pdk1* T-DNA insertion alleles are not likely to be true loss-of-function mutants.

**Figure 2.**
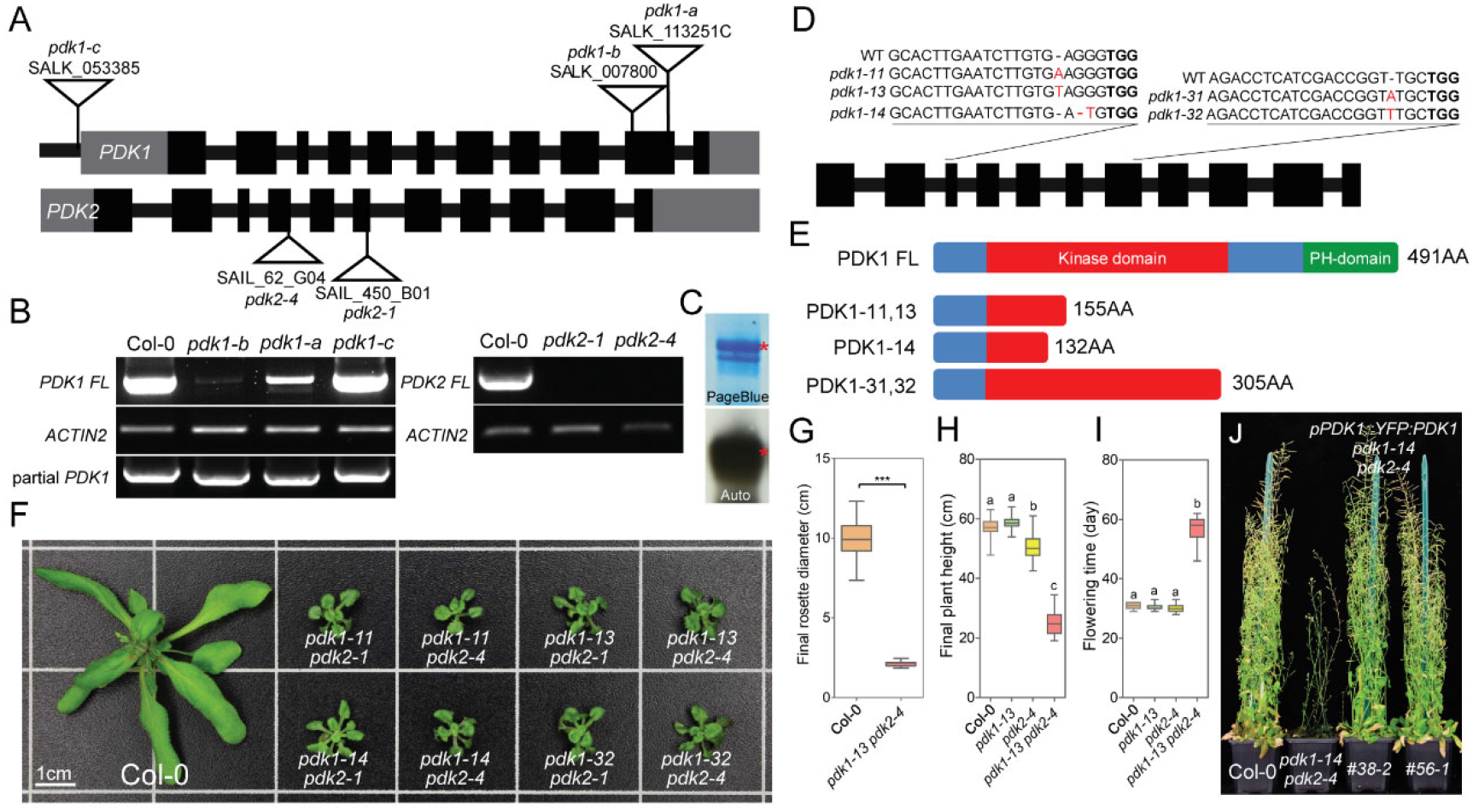
Arabidopsis double mutants that combine the new CRISPR/Cas9-generated *pdk1* loss-of-function alleles with one of the *pdk2* loss-of-function T-DNA alleles exhibit a dwarf stature. A, Schematic representation of the *PDK1* and *PDK2* gene structure, with the T-DNA insertion positions and names of the available mutant alleles indicated. Wide black boxes represent exons, gray boxes represent 5’ and 3’ untranslated regions (UTRs), and narrow black boxes represent introns and promoter sequence upstream of *PDK1* 5’ UTR. B, Semiquantitative RT-PCR to detect *PDK1* or *PDK2* expression in the different *pdk1* or *pdk2* T-DNA insertion mutant alleles, respectively. C, Autophosphorylation activity of GST-PDK1S0 (lacking PH domain, similar size as partial PDK1 in *pdk1-b*. The GST-PDK1S0 band is marked with a red asterisk. D, Schematic representation of part of the *PDK1* gene with the guide RNA target sites and the resulting mutations in the newly obtained CRISPR/Cas9-generated alleles. The PAM sequence for Cas9 is highlighted in bold. Inserted or replaced nucleotides in the new mutant alleles are highlighted in red. Mutant alleles obtained at editing site1 (3^rd^ exon) and site3 (7^th^ exon) are named *pdk1-1n* and *pdk1-3n*, respectively. Since *pdk1-12* and *pdk1-13* have same “T” insertion, only *pdk1-13* is shown. E, Schematic linear representation of the full length (FL, 491 amino acids) PDK1 protein (protein kinase domain: amino acids 44 to 311, PH domain: amino acids 386 to 491, https://www.uniprot.org), and the shorter versions produced in the *pdk1-11, −13, −14, −31* and *-32* alleles. F, Rosette phenotype of 30-day-old wild-type Arabidopsis (Col-0) and eight different *pdk1 pdk2* loss-of-function allele combinations. G, Quantification of the rosette diameter of the 30-day-old Arabidopsis wild-type (Col-0, n=30) and *pdk1-13 pdk2-4* (n=19) plants shown in F. Asterisks indicate significant differences (*t*-test, P<0.0001). H, Final plant height of the indicated lines (n>14). I, Flowering time of the indicated lines (n>14). Letters a, b, and c in H and I indicate statistical differences, as determined by one-way ANOVA followed by Tukey’s test (p<0.05). J, Introduction of *PDK1::YFP:PDK1* in the *pdk1-14 pdk2-4* background completely rescues the mutant phenotype.

In order to obtain true *pdk1* alleles for studying PDK1 biological function, we designed guide RNAs against the 3^rd^ and 7^th^ exon, and were able to obtain five CRISPR/Cas9-generated mutants with frame shifts in the *PDK1* open-reading frame (Figure 2D, E). Like the *pdk1* T-DNA insertion alleles, these new *pdk1* mutant alleles did not result in significant morphological differences from wild type. However, when combined with the *pdk2-1* or *pdk2-4* alleles, all double mutant combinations showed the same striking dwarf phenotype (Figure 2F-J). Complementation analysis using either *p35S::PDK1, p35S::YFP:PDK1* or *pPDK1::YFP:PDK1* showed that the dwarf phenotype was caused by *pdk* loss-of-function (Figure 2J, Figure S2). For all three constructs, several lines were obtained that showed complete rescue of the *pdk1-13 pdk2-4* double mutant phenotype (Figure S2, Figure S3A). The results show that *PDK1* and *PDK2* act redundantly and have a much more important role in plant growth and development than was previously reported.

### *pdk* loss-of-function leads to many developmental defects, but not to a *pid* phenocopy

Besides the decreased rosette diameter and reduced final plant height (Figure 2G, H), *pdk1 pdk2* double mutant plants flowered much later and showed strong reproductive defects (Figure 2I, J; Figure 3). The number of double homozygous F2 progeny obtained was much lower (1 in 47.7 ± 2.6) than the Mendelian ratio (1 in 16). Also F2 plants with the *pdk1*(-/-) *pdk2*(+/-) or *pdk1*(+/-) *pdk2*(-/-) genotype produced homozygous progeny at a much lower frequency than the expected 1 in 4 ratio (Table S1). Seed production of the homozygous *pdk1 pdk2* mutants (1.5 ± 0.21 per silique for *pdk1-13 pdk2-4*) was significantly reduced compared to wild type (65.9 ± 0.61 per silique). Mutant plants developed very short siliques (Figure 3A), a phenotype that has previously been reported for Arabidopsis plants that are both male and female sterile (Huang et al., 2016). These results implied that *pdk1 pdk2* loss-of-function causes gametophyte and/or embryo development defects in Arabidopsis. Reciprocal crosses between wild-type and *pdk1-13 pdk2-4* double mutant plants revealed both male-related and female-related reduced fertility. However, since the cross Col-0♀ x *pdk1-13 pdk2-4*♂ produced fewer seeds than the reciprocal cross *pdk1-13 pdk2-4*♀ x Col-0♂ (Figure S4A), it is likely that male gametophyte development is more strongly impaired by *pdk* loss-of-function than female gametophyte development. Alexander staining showed that pollen grain development in the *pdk1 pdk2* double mutant was not aborted, but that anther dehiscence was the major cause of the male fertility problems (Figure 3B-E, I). In addition, *in vitro* germination of *pdk1 pdk2* pollen resulted in strangely shaped pollen tubes as a result of aberrant tip growth (Figure 3F, G). After 18-hour incubation on pollen germination medium, *pdk1-13 pdk2-4* pollen tube growth arrested with a bulb-like structure, and as a result they remained much shorter than wild type pollen tubes (Figure 3G, H). The ovules of double mutant plants did not show noticeable morphological alterations (Figure S4B,C), which is in line with the predominant effect of *pdk* loss-of-function on male fertility.

**Figure 3.**
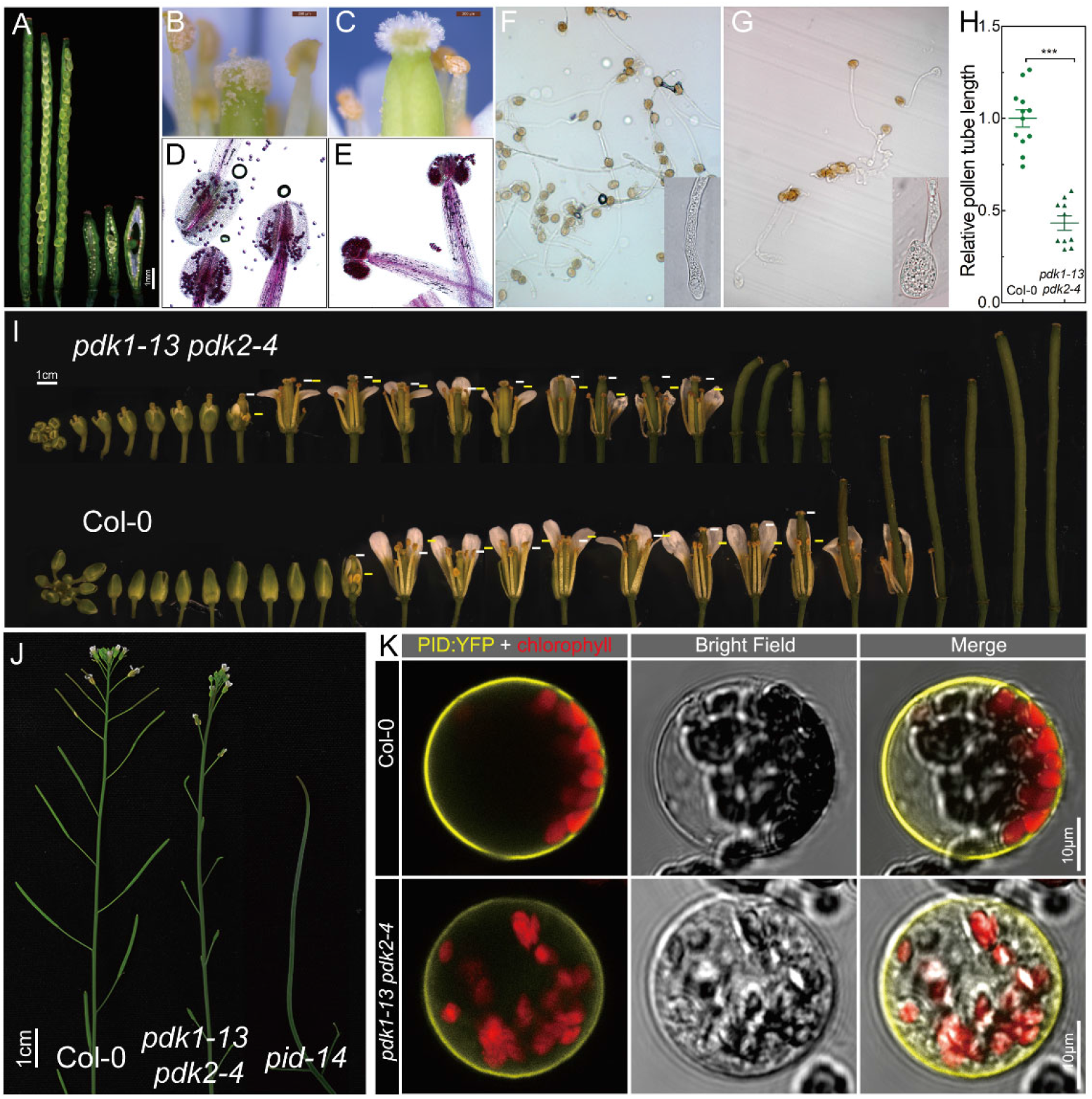
Flowering *pdk1 pdk2* mutant plants show clear developmental defects, but do not phenocopy *pid* mutant plants. A, *pdk1 pdk2* siliques (three on the right) are much shorter than wild-type siliques (three on the left), and contain many unfertilized ovules. B and C, Difference in pollen grain deposition on the stigma of a wild-type (B) or a *pdk1-13 pdk2-4* mutant (C) flower. D and E, Mature wild-type (D) or *pdk1-13 pdk2-4* mutant (E) anthers stained with Alexander’s showing that pollen grains are viable, but that mutant anthers do not sufficiently dehisce. F and G, *In vitro* germination of wild-type (F) and *pdk1-13 pdk2-4* mutant (G) pollen. A detail of pollen tube tip is shown in the inset. H, Relative pollen tube length after 18 hours incubation. The average length of wild-type (Col-0) pollen tubes is put at 1.0. Asterisks indicate a significant difference (Student’s *t*-test, p<0.001). I, Developmental series of *pdk1-13 pdk2-4* mutant and wild-type (Col-0) flowers. The white bars indicates the position of the gynoecium apex, the yellow bars indicate the position of the anthers. J, Inflorescence phenotype of wild type (Col-0), *pdk1-3 pdk2-4* and *pid-14*. K, Representative images of PID:YFP subcellular localization in Col-0 or *pdk1-13 pdk2-4* protoplasts. More than ten observed protoplasts for each line showed the same localization.

In contrast to the fertility problems, *pdk1 pdk2* double mutants developed relatively normal flowers that showed no clear patterning defects. Flowers did show early stigma exposure due to impaired sepal growth, and slightly reduced filament elongation (Figure 3I). The short inflorescences seemed not the result of reduced internode elongation, but were most likely caused by early inflorescence meristem arrest (Hensel et al., 1994) (Figure 2J). The lack of phenotypic resemblance between *pdk1-13 pdk2-4* and *pid-14* inflorescences and flowers (Figure 2J) suggests that PDK1 is not essential for full PID function during inflorescence development. Moreover, expression of a PID:YFP fusion in *pdk1-13 pdk2-4* protoplasts showed that PDK1 activity is not necessary for the predominant localization of PID at the plasma membrane (Figure 2K). Based on these results and the overexpression data we conclude that, in contrast to what has previously been suggested (Zegzouti et al., 2006a, 2006b), PDK1 is not a key regulator of PID activity.

### Alternative splicing produces a functional cytosolic PDK1 isoform lacking a PH domain

The *PDK2* gene produces a single transcript, whereas transcription of *PDK1* results in at least six different mature transcripts, due to alternative splicing events at the 5^th^, 7^th^ and 9^th^ intron (https://www.araport.org/). These transcripts can be translated into five different protein isoforms, which we named respectively PDK1S0, PDK1S1, PDK1S2 and PDK1S3 (Figure 4A). We checked the abundance of each mature transcript using semi-quantitative RT-PCR followed by restriction digestion. The full-length *PDK1* transcript was most abundant, and the short *PDK1* transcripts producing isoforms lacking part of the kinase domain (PDK1S1, PDK1S2, and PDK1S3) were also present at high levels, while the transcript producing the PDK1S0 isoform with a complete kinase domain, was the least abundant (Figure 4B).

**Figure 4.**
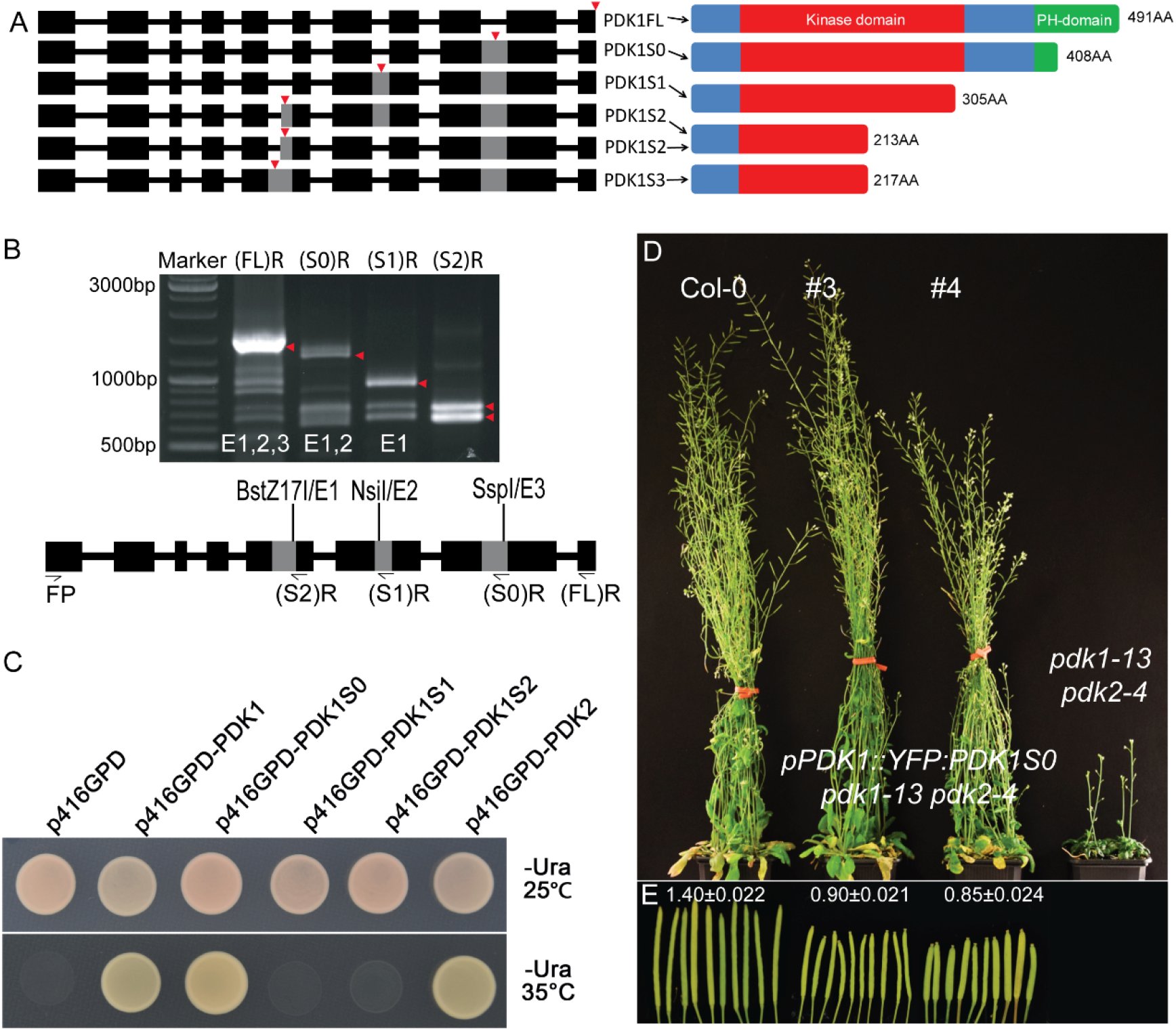
Alternative splicing produces a *PDK1* transcript encoding a functional PDK1 variant lacking the PH domain. A, Schematic representation of the *PDK1* gene indicating the alternative splice events (left) and the respective protein isoforms produced by the splice variants (right). On the left: wide black boxes represent exons, gray boxes represent unspliced introns, black lines represent spliced introns, red arrows point out the locations of stop codons. Please note that PDK1S1 is lacking six amino acids of the kinase domain. B, Expression level of the different splice variants, as detected by RT-PCR. Primer binding and restriction enzyme recognition site locations are shown in the schematic representation below (see detailed description in the materials and methods section). Red arrowheads point out the *PDK1FL*, *PDK1S0*, *PDK1S1* and *PDK1S2/3* transcripts (from left to right) detected using the reverse primers and restriction enzymes indicated above and below the gel image, respectively. C, Rescue of the temperature-sensitive growth of the yeast *pkh1 pkh2* mutant strain by expression of the *PDK1* or *PDK2* full length cDNA or the *PDK1S0* splice variant cDNA. Three biological repeats showed the same result. D and E, The *pPDK1::YFP:PDK1S0* construct rescues the delay in flowering time, short plant height and dwarf rosette leaves of the *pdk1-13 pdk2-4* mutant (D, 14 independent lines were observed), but plants develop shorter siliques carrying less seeds (E). Shown in (D) are 65-day-old plants. Numbers above siliques in (E) represent average length ± SEM (n=10).

In order to test the functionality of the different isoforms, we expressed the corresponding cDNAs in yeast (*S. cerevisiae*) strain INA106-3B, in which the *PKH2* gene copy has been replaced by *LEU2*, and the *PKH1* gene copy has been mutated so that strain INA106-3B is able to grow normally at 25°C but not at 35°C. As expected based on previous experiments, expression of the full length *PDK1* or *PDK2* cDNAs allowed this strain to grow at 35°C (Dittrich and Devarenne, 2012a) (Figure 4C). In contrast, expression of the cDNAs producing the PDK1S1, PDK1S2 or PDK1S3 isoforms did not allow growth at the restrictive temperature, suggesting that any deletion of the conserved kinase domain renders PDK1 non-functional (Figure 4C). This is in line with loss-of-function observed for the new *Arabidopsis* alleles *pdk1-11, −13, −14, −31* and -*32*, which all express partial PDK1 proteins having a small or bigger deletion of the C-terminal part of the kinase domain (Figure 2D, E, F). Interestingly, expression of the PDK1S0 did permit INA106-3B to grow at 35°C (Figure 4C). The yeast data were confirmed by *35S* promoter-driven expression in the Arabidopsis *pdk1 pdk2* loss-of-function mutant background. *p35S::PDK1* provided full rescue of the vegetative growth phenotypes of the Arabidopsis *pdk1 pdk2* mutant, and some *p35S::PDK1S0* lines showed the same level of rescue (Figure S3A). In contrast, expression of PDK1S1 and PDK1S2 did not result in any rescue (Figure S3A). Expression of a *YFP:PDK1S0* fusion under control of the *PDK1* promoter in the *pdk1-14 pdk2-4* mutant background also completely rescued the mutant vegetative growth phenotypes (Figure 4D). However, *pPDK1::YFP:PDK1S0 pdk1-13 pdk2-4* plants developed shorter siliques carrying fewer seeds compared to wild-type or *pPDK1::YFP:PDK1 pdk1-14 pdk2-4* plants (Figure 4E). Interestingly, a similar silique phenotype has also been described for the Arabidopsis *pdk1-b pdk2-1* double mutant, and according to our own analysis the T-DNA insertion in the *pdk1-b* allele leads to the production of a shorter PDK1 protein with an intact kinase domain but lacking the PH domain (Camehl et al., 2011) (Figure 2A, B).

These results corroborate the conclusions from the complementation experiments in yeast that a full-length kinase domain is essential for PDK1 function, but that surprisingly the PH domain is not essential for PDK1 function during Arabidopsis vegetative growth. Since the PH domain is responsible for lipid binding, we checked the *PDK1* promoter driven YFP:PDK1 and YFP:PDK1S0 localization in root columella cells, where *PDK1* is highly expressed. YFP:PDK1 localized both on the plasma membrane and in the cytoplasm, whereas YFP:PDK1S0 was only found in the cytoplasm (Figure S3B, C). The functional relevance of the latter low abundant cytosolic isoform remains unclear. Our findings do suggest, however, that PH domain-dependent plasma membrane association of PDK1 is only essential during specific developmental processes.

### *PDK1* and *PDK2* are broadly expressed during development

Since the *pdk1 pdk2* mutant shows many defects in development and growth, we analysed the spatio-temporal expression pattern of the two *PDK* genes to uncover their tissue-specific functions. For this purpose we generated Arabidopsis (Col-0) lines carrying the *pPDK1::turboGFP:GUS (pPDK1-GG)* or *pPDK2::turboGFP:GUS (pPDK2-GG)* construct and used a *pdk1-14 pdk2-4* mutant line carrying the complementing *pPDK1::YFP:PDK1* construct. *PDK1* appeared to be strongly expressed in (pro)vascular tissues from the early globular embryo stage on, and in the columella root cap (Figure 5, Figure S5). The gene also showed more general expression in young hypocotyls, cotyledons, leaves and floral organs, and in growing siliques. The expression pattern of *PDK2* was very comparable to that of *PDK1* (Figure 5 D, E, H, K, L), except that no expression was observed in the root apex (Figure 5D) or in embryos (data not shown).

**Figure 5.**
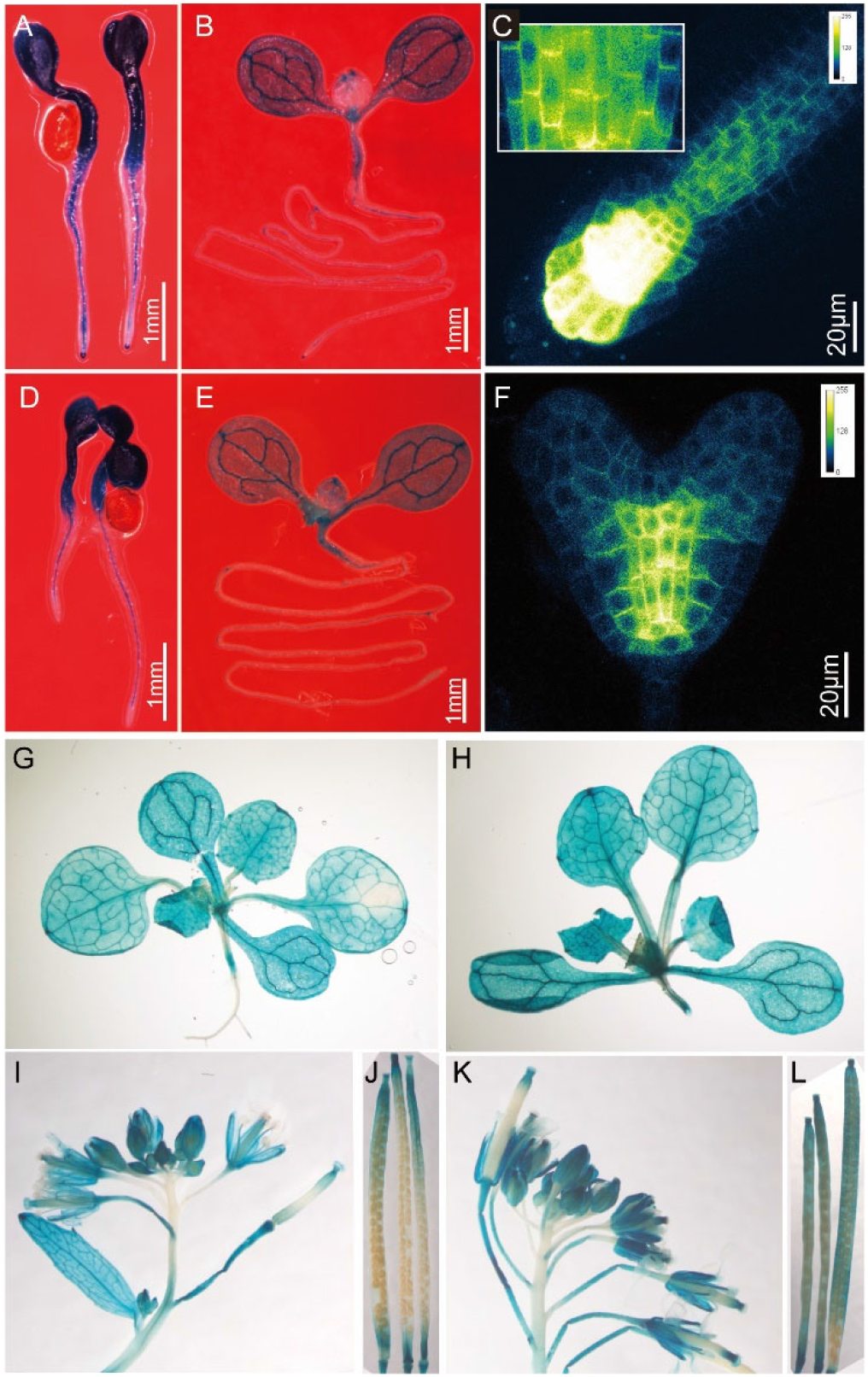
*PDK1* and *PDK2* are predominantly expressed in (pro) vascular tissues, where PDK1 associates with the basal plasma membrane. Spatio-temporal expression pattern of *PDK1* (A, B, G, I, J) and *PDK2* (D, E, H, K, L) as reported by histochemical staining of 3-day-old seedlings (A, D), 7-day-old seedlings (B, E), 16-day-old plants (G, H) and inflorescences and siliques from 40-day-old plants (I-L) of representative *pPDK1-GG* and *pPDK2-GG* lines, respectively. C and F, Confocal images of a 4-day-old *pPDK1::YFP:PDK1* root tip (C) or a heart stage embryo of the same transgenic line (F). The inset in (C) shows a detail of YFP:PDK1 localization in root stele cells.

As previously observed for PDK2:EGFP (Scholz et al., 2019), YFP:PDK1 did not localize in the nucleus, but was mainly found in the cytoplasm or associated with the plasma membrane (Figure 5 C, F; Figure S3B;). In (pro)vascular cells in heart stage embryos and roots, YFP:PDK1 showed predominant basal (rootward) localization (Figure 5 C, F; Figure S5), just like PIN1 and the PDK1 targets PAX and D6PK (Gälweiler et al., 1998; Zourelidou et al., 2009; Marhava et al., 2018).

### Auxin transport is impaired in *pdk1-13 pdk2-4* mutant

Even though the *PDK1* overexpression and loss-of-function phenotypes did not point to an important role for PDK1 in PID function, the *pdk1-13 pdk2-4* mutant seedling phenotypes did suggest involvement of PDK1 in the regulation of auxin response or -transport (Figure 6A-E). Mutant primary roots elongated normally up to two days after germination, but after that their growth rate declined (Figure 6 A,B), and roots started to oscillate randomly with a large amplitude, resulting in curved short roots (Figure S6). Of 199 7-day-old seedlings, 18.1% of the primary roots grew into the air. On a total of 460 *pdk1-13 pdk2-4* mutant seedlings, 58% showed fused or single dark green cotyledons, and the remaining 42% developed two cotyledons with short petioles positioned at an abnormal angle (< 180°) (Figure 6C-E). The cotyledon phenotypes and short agravitropic roots are usually observed in auxin response or -transport mutants or transport inhibitor-treated seedlings. By combining the *pdk1-13 pdk2-4* double mutant with the or *pDR5::GUS* auxin response reporter, we observed that auxin response was absent or strongly decreased in the root stele and confined to the root tip, while an enhanced auxin response was observed at the mutant cotyledon edges and in the fragmented cotyledon veins (Figure 6F, G, I, K). This highly resembled the *DR5::GUS* expression of 7-day-old seedlings grown on medium supplemented with the auxin transport inhibitor naphthylphtalamic acid (NPA) (Figure 6H, J) (Sabatini et al., 1999; Bao et al., 2004). Moreover, the increase in *DR5::GUS* expression in cotyledons corroborated that *pdk1-13 pdk2-4* mutants are defective in auxin transport, rather than in auxin biosynthesis or -signaling. Short time treatment of wild-type and *pdk1-13 pdk2-4* seedlings with IAA and subsequent qPCR analysis showed that the auxin inducible expression of the *IAA5*, *GH3.3* and *SAUR16* genes was not impaired, confirming that the mutants are not defective in auxin response (Figure 6L). Instead, *pdk1-13 pdk2-4* mutant seedlings were hypersensitive to NPA treatment compared to wild type (Figure 6M). Moreover, the auxin transport capability of *pdk1-13 pdk2-4* inflorescence stems was significantly reduced compared to that of wild-type stems (Figure 6N). Together, the above data point toward a role for PDK1 in enhancing polar auxin transport.

**Figure 6.**
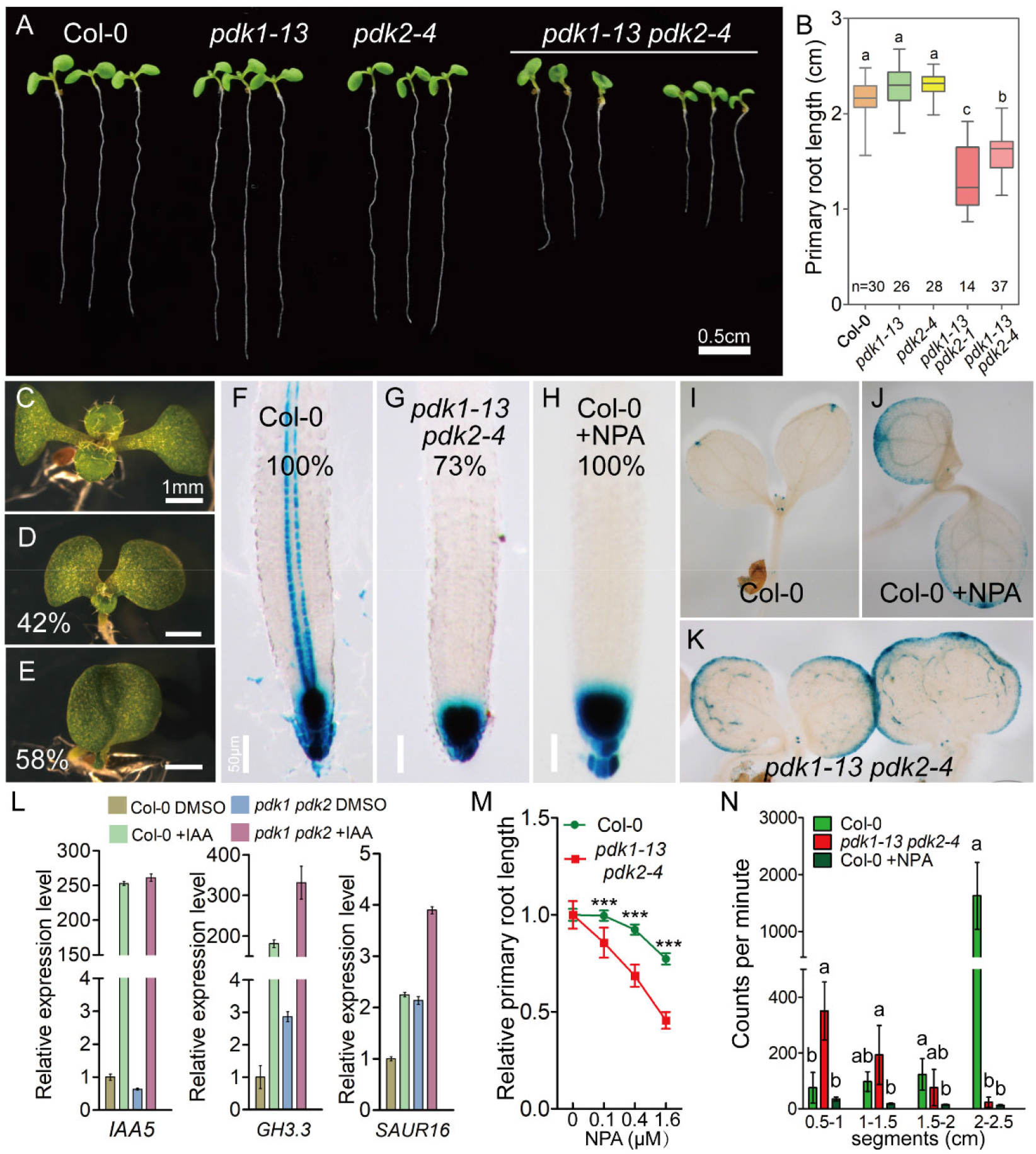
The phenotype and auxin response pattern of *pdk1pdk2* mutant seedlings resembles that of auxin transport inhibitor treated seedlings. A, The phenotype of aligned 7-day-old wild-type (Col-0) and *pdk1 pdk2* mutant seedlings. B, Primary root length of 7-day-old seedlings. Lowercase letters indicate averages that are significantly different, as tested by a one-way ANOVA followed by Tukey’s test (p<0.05). C-E, Cotyledon phenotype of wild-type (Col-0) seedlings (C), showing two symmetrically distributed cotyledons with extended petioles, or *pdk1-13 pdk2-4* seedlings, of which 42% position at an angle with short petioles (D, n=460) and 58% show fused cotyledons (E, n=460). F-K, Histochemical GUS staining of 7 day old wild-type seedlings (Col-0, F, I), wild-type seedlings grown on 0.5μM NPA (Col-0 + NPA, H, J), or *pdk1-13 pdk2-4* seedlings (G, K), all three containing the *pDR5::GUS* auxin response reporter. Percentages in F-H indicate the ratio of representative image out of the observed seedlings (n=15). The rest 27% of *pdk1-13 pdk2-4* show strongly decreased but not absent *pDR5::GUS* signal in the stele. L, Quantitative RT-PCR analysis of the auxin-induced expression of *IAA5*, *GH3.3* and *SAUR16* in 5-day-old Arabidopsis wild-type (Col-0) and *pdk1-13 pdk2-4* mutant seedlings. The values displayed in the graph are means ± SEMs. M, NPA sensitivity of wild-type (Col-0) and *pdk1-13 pdk2-4* based on the primary root length of seedlings grown on medium with an increasing NPA concentration (n>22, Student’s *t*-test was used for analysis between groups from the same NPA concentration, p<0.001). Error bar = 95% confidence interval. N, Transport of ^3^H-IAA in 2.5 cm wild-type (Col-0), wild-type with NPA and *pdk1 pdk2* inflorescence stem pieces. Bars represent the average number of counts per segment ± 95% confidence interval. Data were analyzed using a one-way ANOVA followed by Tukey’s test. Significant differences are indicated with different letters in each segment group. A representative experiment of three biological repeats displaying similar results is shown.

### PDK1 regulates PIN-mediated auxin efflux probably through the AGC1 kinases

Several *in vitro* PDK1 phosphorylation substrates are AGC kinases that have been reported to regulate auxin transport by direct phosphorylation of PIN auxin efflux carriers (Zegzouti et al., 2006b; Willige et al., 2013; Marhava et al., 2018; Haga et al., 2018). The fact that their loss-of-function mutants share phenotypic defects with the *pdk1-13 pdk2-4* mutant, such as the short root of the *pax* mutant (Marhava et al., 2018), or the fused cotyledons of the *d6pk012* triple mutant (Zourelidou et al., 2009), hinted that these AGC kinases might indeed be regulated by PDK1 *in planta*. The stronger *pdk1-13 pdk2-4* mutant phenotype suggested that PDK1 has many more phosphorylation substrates.

In order to investigate whether PIN proteins themselves are PDK1 phosphorylation substrates, we deduced based on published *in vitro* phosphorylation data, that PDK1 prefers to phosphorylate the second serine residue in the RSX<u>S</u>FVG motif (X represents any amino acids) that is part of the activation segment of the AGC kinases (Zegzouti et al., 2006a, 2006b). Analysis of the large central hydrophilic loop (HL) of the 5 Arabidopsis PIN1-type PIN proteins identified several RXXS motifs. However, *in vitro* phosphorylation assays using GST-tagged PDK1 (GST-PDK1) and GST-tagged versions of the HL of PIN1, PIN2, PIN3 or PIN7 (GST-PIN1/2/3/7HL) only showed phosphorylation of the PIN2HL (Figure 7A). Interestingly, PIN2HL S1,2,3A, in which the PID phosphorylation sites are substituted by alanines, was also phosphorylated by PDK1 at same level as the wild-type PIN2HL (Figure 7A). PDK1 must therefore phosphorylate one or more other serine residues that are unique to the PIN2HL. However, PIN2 is not co-expressed with PDK1, and the PIN proteins that are co-expressed with PDK1 in the root stele or columella cells (PIN1, PIN3 and PIN7) are not phosphorylated by PDK1 *in vitro*. Moreover, no noticeable alteration in PIN1/3/7 protein polarity was observed in *pdk1-13 pdk2-4* mutant roots (Figure 7C-J). The PIN2:GFP abundance was slightly decreased in *pdk1-13 pdk2-4* mutant root tips (Figure 7I, J, L), but this might be an indirect effect of *pdk* loss-of-function on auxin distribution in the root tip, as we measured a slight increase in GFP intensity in *DR5::GFP pdk1-13 pdk2-4* mutant versus wild-type root tips (Figure 7B, K). Our results suggest that PDK1 regulates auxin transport, most likely by activating one or more AGC kinases, such as PAX and D6PK, which subsequently regulate auxin efflux activity by direct phosphorylation of the PINs.

**Figure 7.**
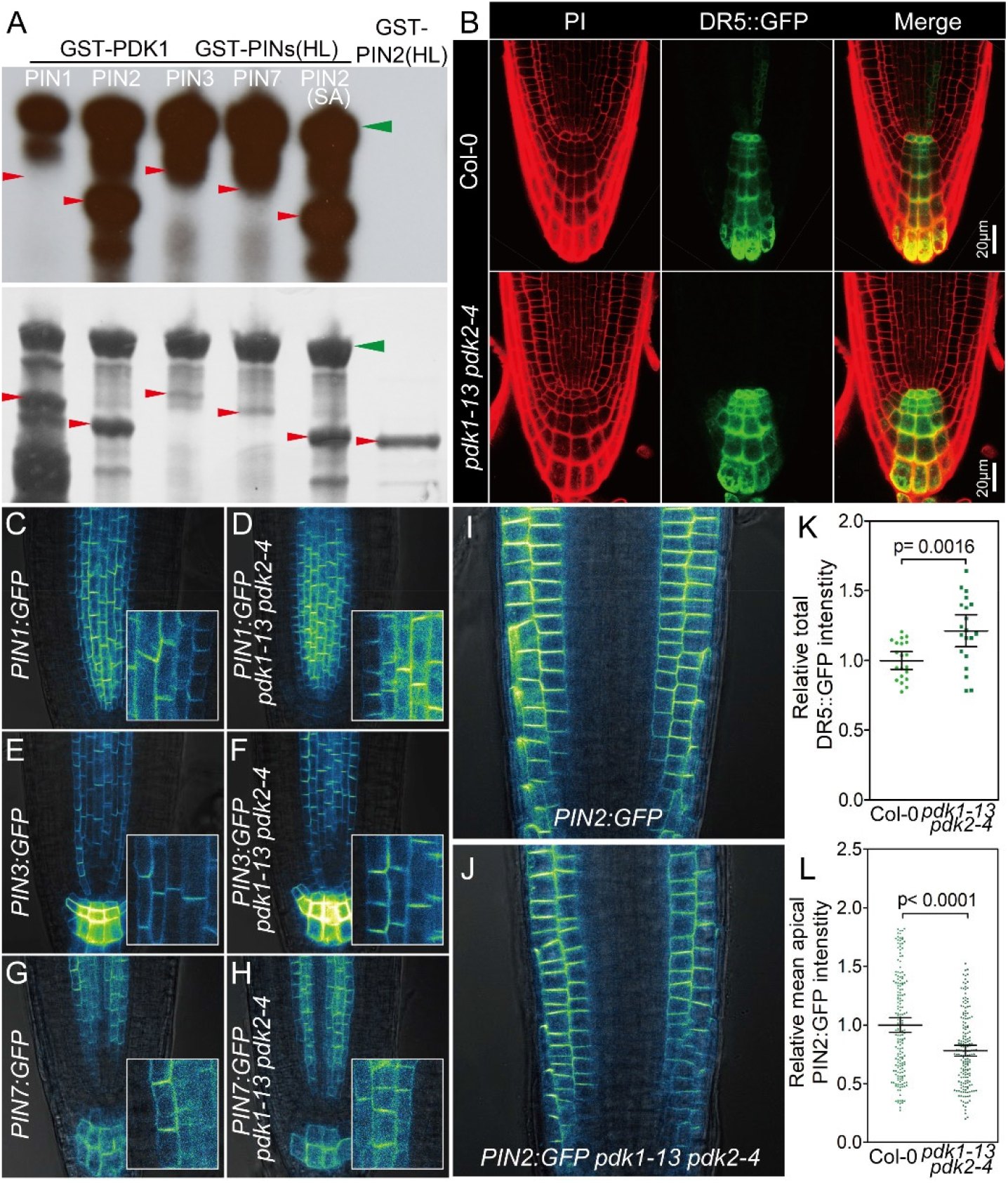
PDK1 is not involved in PIN polarity control. A, PDK1 phosphorylates the PIN2HL, but not the PIN1HL, PIN3HL or PIN7HL, in a S1, S2, S3-independent manner *in vitro*. Red or green arrows point out the position of the GST-PINHL or GST-PDK1, respectively. Upper: autoradiograph, lower: PageBlue stained gel. B, Confocal images of *DR5::GFP* expression in a wild-type (Col-0, upper panel) or a *pdk1-13 pdk2-4* mutant root tip (lower panel). Left: propidium iodide (PI) staining; middle: GFP signal; right: merged image. C-J, Confocal images showing the subcellular localization of PIN1:GFP (C, D), PIN3:GFP (E, F), PIN7:GFP (G, H), and PIN2:GFP (I, J) in wild-type (Col-0, C, E, G, I) or *pdk1-13 pdk2-4* mutant (D, F, H, J) root tips. Insets in C-H show the details of PIN polarity in stele cells. K and L, Relative total GFP intensity produced by *DR5::GFP* expression in columella-QC cells (K, n=20) or representing PIN2:GFP at the apical side of the epidermal cells (L, n=8), in wild-type (Col) or *pdk1-13 pdk2-4* mutant roots. DR5::GFP and PIN2:GFP intensities are shown relative to Col-0 control.

## Discussion

PDK1 is a well-established key regulator of AGC kinases in animals and yeast, and its importance in these organisms is demonstrated by the lethality caused by loss-of-function mutations in the genes encoding for this protein kinase (Casamayor et al., 1999; Rintelen et al., 2002; Lawlor et al., 2002; Mora et al., 2004). Also in the model plant Arabidopsis, PDK1 has been shown to phosphorylate several AGC kinases *in vitro* (Zegzouti et al., 2006a, 2006b), However, the previously reported impact of loss-of-function of the two gene copies *PDK1* and *PDK2* on Arabidopsis development was only limited (Camehl et al., 2011; Scholz et al., 2019). In this study, we found that the published T-DNA insertion alleles of the Arabidopsis *PDK1* gene copy are not loss-of-function mutants. Here we generated several CRISPR/Cas9-based true loss-of-function *pdk1* alleles that, when combined with the available *pdk2* loss-of-function mutant alleles, did lead to strong developmental defects. Different from animals and yeast though, and more similar to the situation in *Physcomitrella Patens*, Arabidopsis *pdk1 pdk2* loss-of-function mutants are viable, indicating that the substrate preference of plant PDK1 has changed from that in other eukaryotes, and that it has lost its involvement in signaling pathways that are essential for cell survival.

### PDK1 is not essential for PID activity controlling inflorescence and cotyledon development

By carefully recording the *pdk1 pdk2* mutant phenotypes, we analysed the genetic relation between PDK1 and its reported *in vitro* substrates, of which PID was the key candidate (Zegzouti et al., 2006a). Loss-of-function of both *PDK* genes leads to fused cotyledons, short wavy roots, dwarf stature and reduced fertility resulting in short siliques. To our surprise, *pdk1 pdk2* does not share the three cotyledon, pin inflorescence and aberrant flower phenotypes that are typical for *pid* loss-of-function mutants, implying that PDK1 is not essential for PID function in these tissues. PID is an auto-activating kinase *in vitro* and might act independent of upstream activating kinases (Christensen et al., 2000; Benjamins et al., 2001), or other kinases than PDK1 might be involved in hyper-activating PID during embryo, inflorescence and flower development. The latter seems most likely based on the observation that flower, leaf and shoot extracts can hyperactivate PID *in vitro* (Zegzouti et al., 2006a). A physical interaction between PID and PDK1 through the PIF domain, as suggested by Zegzouti et al. (Zegzouti et al., 2006a), has never been proven, and was purely based on *in vitro* phosphorylation data. Here we show unequivocally that PID does not require PDK1 for its association with the PM, which corroborates the finding that this is mediated by an arginine-rich loop in the kinase domain of PID (Simon et al., 2016). All data are in line with the observation that a PID:GUS fusion lacking the PIF domain can still complement *pid* loss-of-function mutants (Benjamins et al., 2001). Although we cannot fully exclude that PDK1 and PID do have a functional interaction, our results at least indicate that this interaction is not essential for the majority of the PID activities in plant development.

### Is PDK1 a master regulator of AGC kinases in Arabidopsis?

If not PID, which and how many other Arabidopsis AGC kinases are potential phosphorylation substrates of PDK1? The developmental defects of the *pdk1 pdk2* double mutant together with the altered *pDR5* expression pattern and reduction in auxin transport all point toward a role for PDK1 in promoting polar auxin transport. Interestingly, several *pdk1 pdk2* mutant phenotypes are also observed for loss-of-function mutants of members of the AGC1 kinase sub-family, for some of which a role as regulator of polar auxin transport is now well established (Zourelidou et al., 2009; Willige et al., 2013; Barbosa et al., 2014; Haga et al., 2018; Marhava et al., 2018). For example, the fused cotyledons, deficient lateral root emergence and agravitropic primary root growth closely resemble phenotypes observed for the *d6pk012* triple mutant (Zourelidou et al., 2009). And a short primary root is also observed for the *pax* mutant (Marhava et al., 2018), and like *pdk1 pdk2*, the *agc1.5 agc1.*7 double mutant is defective in pollen tube growth (Zhang et al., 2009). Since the corresponding AGC1 kinases are strongly dependent on PDK1 for their *in vitro* activation (Zegzouti et al., 2006b), it seems possible that PDK1 might act as a master regulator of these AGC1 kinases. Further experimentation is required, however, for each of these kinases to prove this hypothesis.

Recently, a genetic interaction with *PDK1* was reported for a kinase of the AGC2 clade, UNICORN (UCN, AGC2-3), in controlling integument growth (Scholz et al., 2019). According to Scholz and coworkers, UCN acts as repressor of PDK1 function. The absence of UNC activity or PDK1 overexpression leads to uncontrolled integument and petal growth, and the *pdk1-c pdk2* T-DNA insertion mutant combination can rescue the *ucn* loss-of-function mutant defects. Like Scholtz et al., we did not observe defects in ovule development for the new *pdk1 pdk2* allelic combinations (Figure S4B, C). It will be interesting to see if our new *pdk1-13 pdk2-4* double mutant combination will also lead to restoration of the *ucn-1* flower and ovule phenotypes.

### Alternative splicing provides possible functional differentiation for PDK1

By studying *pdk1* T-DNA insertion alleles and splice variants produced by the *PDK1* gene, we revealed that the PH domain is not essential for the general PDK1 function in Arabidopsis. The alternative splicing product PDK1S0, which lacks phospholipid binding ability and membrane localization but still has kinase activity, is able to rescue the thermosensitive growth of the yeast *pkh1 pkh2* double mutant strain INA106-3B, and most of the developmental defects of the *Arabidopsis pdk1 pdk2* loss-of-function mutant. A similar PDK1 protein variant appeared to be produced in the Arabidopsis *pdk1-a* and *pdk1-b* T-DNA insertion alleles, which were initially thought to be complete loss-of-function alleles. This explains the relatively mild flower phenotypes observed for the Arabidopsis *pdk1-a pdk2* and *pdk1-b pdk2* double mutants (Camehl et al., 2011; Scholz et al., 2019). The reduced growth response of these mutants induced by phosphatidic acid downstream of *Piriformospora indica* infection (Camehl et al., 2011) suggests a differential function for PDK1S0 and PDK1 in development and stress response, respectively. This is in line with the observation that phosphatidic acid, an important second messengers for stress response, can directly bind and stimulate the activity of full length Arabidopsis PDK1 (Deak et al., 1999; Anthony et al., 2004). In animals, phospholipids are known to bind to PDK1 to induce PDK1 dimer to monomer conversion and activation (Alessi et al., 1997; Ziemba et al., 2013). The functionality of PDK1S0 in most developmental processes questions the importance of the clear basal polarity of full length PDK1 in (pro) vascular cells in the embryo and root tip. Apparently, PDK1 binding to the AGC kinase PIF domain is sufficiently efficient, and does not require prior co-localisation at the PM. In conclusion, alternative splicing of *PDK1* transcripts may provide a novel and unique regulation mechanism for balancing growth and defense in Arabidopsis, which differs from animals and yeast.

## Materials and methods

### Plant lines and growth condition

*Arabidopsis thaliana (L.)* ecotype Columbia 0 (Col-0) was used as wild-type control for all experiments, since all mutant and transgenic lines are in the Col-0 background. Previously described T-DNA insertion lines SALK_053385 (*pdk1.1-1,* renamed to *pdk1-c*), SALK_11325C (*pdk1.1a*, renamed to *pdk1-a*), SALK_007800 (*pdk1.1b*, renamed to *pdk1-b*), SAIL_62_G0**<u>4</u>** (*pdk1.2-2*, renamed to *pdk2-4*) and SAIL_450_B0**<u>1</u>** (*pdk1.2-3*, renamed to *pdk2-1*) were ordered from the Nottingham Arabidopsis Stock Centre (Camehl et al., 2011; Scholz et al., 2019). The following Arabidopsis lines are also described elsewhere: *pDR5::GFP* (Ottenschlager et al., 2003), *pDR5::GUS* (Benjamins et al., 2001), *pPIN1::PIN1:GFP* (Benkova et al., 2003), *pPIN2::PIN2:GFP* (Xu and Scheres, 2005), *pPIN3::PIN3:GFP* (Zádníková et al., 2010), *pPIN7::PIN7:GFP* (Blilou et al., 2005) and *p35S::PID#21* (Benjamins et al., 2001). For lines created in this study, the T-DNA constructs *p35S::YFP:PDK1*, *p35S::PDK1*, *pPDK1/2::turboGFP:GUS,* and *pYAO-Cas9-gRNA1/2/3* were transformed into Col-0 using *Agrobacterium*-mediated floral dip transformation (Clough and Bent, 1998). Homozygous lines with a single T-DNA insertion were selected for further analysis. Of the 80 CRISPR/Cas9 transgenetic alleles obtained, 7 appeared to contain loss-of-function mutations in the 3^rd^ and 7^th^ exon of *PDK1*. The CRISPR/Cas9 T-DNA construct in the new *pdk1* mutant alleles was segregated out during the generation of the *pdk1 pdk2* double mutant. Five mutant alleles with open reading frame shifts were used for further analysis (Figure 2C).

For complementation analysis of PDK1 isoforms, the T-DNA constructs *p35S::YFP:PDK1* or *p35S::PDK1FL/S1/S2* were transformed into the *pdk1-13(+/-) pdk2-4(-/-)* mutant background, *p35S::YFP:PDKS0* or *p35S::PDK1S0* were transformed into the *pdk1-13(-/-) pdk2-1(+/-)* mutant background, or *pPDK1::YFP:PDK1FL and S0* were transformed into the *pdk1-14(-/-) pdk2-4(+/-)* or *pdk1-13(+/-) pdk2-4(-/-)* mutant background, respectively. The genotype of the *pdk1 pdk2* mutant background was confirmed by PCR before floral dip transformation. All genotyping primers are summarized in Table S2.

Plants were grown on soil at 21 ℃, 16 hr photoperiod, and 70% relative humidity. For seedling growth, seeds were surface-sterilized by 1 minute in 70% ethanol, 10 minutes in 1% chlorine followed by five washes with sterile water. Sterilized seeds were kept in the dark at 4 °C for 2 days for vernalization and germinated on vertical plates with 0.5× Murashige and Skoog (1/2 MS) medium (Duchefa) containing 0.05% MES, 0.8% agar and 1% sucrose at 22 °C and 16 hr photoperiod.

### RNA extraction and (q)RT-PCR

Total RNA was extracted from 5-day-old seedlings using a NucleoSpin RNA Plant kit (Macherey Nagel, #740949). Reverse transcription (RT) was performed using a RevertAid Reverse Transcription Kit (Thermo Scientific™, #K1691). For qRT-PCR on auxin induced genes, RNA was isolated from 5-day-old Col-0 and *pdk1-13 pdk2-4* seedlings treated for 1 hour with 10 μM IAA. Gene expression was normalized to the reference gene *PP2A-3* (*AT2G42500*) using the ΔΔCt method. For analysis of the *pdk1* and *pdk2* T-DNA alleles, RT-PCR was performed with DreamTaq DNA Polymerases (Thermo Scientific™). (q)RT-PCR primers are listed in Table S2. For detection of the *PDK1* splice variants, RT-PCR was performed for 40 cycles using the forward (FP) and reverse (FL, S0, S1, and S2) primers (Figure 5B), as listed in Table S2. PCR reactions with primer pair FP and (FL)R were digested with *Bst*Z17I, *Nsi*I and *Ssp*I to detect PDK1FL, with primer pair FP and (S0)R with *Bst*Z17I and *Nsi*I to detect PDK1S0, and with primer pair FP and (S1)R with *Bst*Z17I to detect PDK1S1. 0.1 μL of the enzymes *Bst*Z17I, *Nsi*I and/or *Ssp*I (Thermo Scientific™) was directly added to the 20 μL PCR reaction and reactions were incubated at 37 ℃ overnight before gel electrophoresis. Detection of PDK1S2 and PDK1S3 with primer pair FP and (S2)R did not require restriction enzymes digestion. qRT-PCR was performed in the CFX96 Touch™ Real-Time PCR Detection System (Bio-Rad) using TB Green Premix Ex Taq II (Tli RNase H Plus) (Takara, #RR820B).

### Cloning procedures

To generate the *Promoter::turboGFP:GUS* fusions, a *Sac*I-*TurboGFP*-*Pac*I fragment was cloned from *pICSL80005* into *pMDC163*, resulting in *pMDC163(gateway)-TurboGFP:GUS*. *PDK1* and *PDK2* promoter regions of approximately 2.0 kb including the first six codons were amplified from Col-0 genomic DNA using the primers listed in Table S2, and cloned in *pDONR207* by LR recombination. The resulting fragments were subsequently fused in-frame with the *turboGFP*:*gusA* reporter gene in *pMDC163(gateway)-TurboGFP:GUS* by BP recombination. (Invitrogen, Gateway BP/LR Clonase II Enzyme Mix, #11789020 and #12538120).

*PDK1* splice variants were amplified from cDNA of 5-day-old seedlings using the respect primers (Table S2), after which restriction enzymes described in the RT-PCR section were employed. Fragments were cloned in *pDONR207* by BP recombination, and subsequently transferred to *pART7-35S::YFP:gateway* by LR recombination, resulting in *pART7-35S::YFP:PDK1FL/S0/S1/S2*. Expression cassettes were excised with *Not*I and cloned into *Not*I digested *pART27*, resulting in *pART27-35S::YFP:PDK1FL/S0*. The same entry vectors and LR recombination were used to generate *pMDC32-35S::PDK1FL/S0/S1/S2* and *pGEX-PDK1FL/S0*. *pGEX-PIN1HL*, *pGEX-PIN2HL* and *p35S:PID:YFP* have been described previously (Galván-Ampudia and Offringa, 2007; Huang et al., 2010; Dhonukshe et al., 2010). *PIN3HL* and *PIN7HL* were amplified from Col-0 cDNA using primers listed in Table S2 and cloned into *pGEX* also using Gateway cloning technology to obtain *pGEX-PIN3HL* and *pGEX-PIN7HL*.

To generate *pPDK1::YFP:PDK1FL/S0* fusions, the 2.0Kb *PDK1* promoter region was introduced into *pART27-35S::YFP:PDK1S0* by replacing the *35S* sequence using restriction enzymes *Bst*XI and *Kpn*I. *pART27-pPDK1::YFP:PDK1S0* and *pDONR207* were mixed with BP clonase to obtain *pART27-pPDK1::YFP:gateway*. *PDK1FL* was then recombined into *pART27-pPDK1::YFP:gateway* by LR reaction to obtain *pART27- pPDK1::YFP:PDK1FL*.

To obtain the *p416GPD-PDK* constructs for expression in yeast, *Bam*HI-*PDK1FL/S0/S1/S2*-*Eco*RI and *Bam*HI-*PDK2*-*Xho*I fragments were amplified from *pDONR207-PDK1FL/S0/S1/S2* and 5-day-old seedling cDNA, respectively, using primers listed in Table S2. Fragments were digested with the appropriate restriction enzymes and ligated into vector *p416GPD*.

The *pCambia-pYAO-Cas9-gRNA1/2/3* plasmids for CRISPR/Cas9 mediated mutagenesis were obtained by ligating the *Eco*RI-(Cas9+terminator)-*Avr*II fragment from pDE-Cas9 (Fauser et al., 2014) into *pCambia1300* digested with *Eco*RI and *Xba*I. The *Eco*RI and *Sal*I sites in the resulting *pCambia-Cas9* plasmid were used to clone the *Eco*RI*-YAO* promoter*-Eco*RI (Yan et al., 2015) and *Xho*I-gateway-*Xho*I fragments amplified from respectively *Arabidopsis* Col-0 genomic DNA and the *pART7-35S::YFP:gateway* plasmid. Regions producing guide RNAs (Table S2) designed to target respectively the 3^rd^, 6^th^ or 7^th^ exon of *PDK1* were ligated into pEn-Chimera (Fauser et al., 2014), and introduced behind the *YAO* promoter in pCambia-pYAO-Cas9-gateway by LR recombination.

All primers used for cloning are summarized in Table S2.

### General phenotypic analysis and physiological experiments

NPA treated (stock in DMSO,1/10^4^ dilution) or normally grown seedlings, potted plants, siliques and inflorescences were photographed with a Nikon D5300 camera at the indicated time. For imaging of inflorescences, the top part of the inflorescence was cut from 15 cm high plants. For Figure 6A, seedlings were transferred to and aligned on a black plate before imaging. Primary root length, rosette diameter and silique length were measured with ImageJ (Fiji). Plant height was measured directly using a ruler. Root tips, opened siliques, flowers, details of floral organs and cotyledons were imaged using a Leica MZ16FA stereomicroscope equipped a with Leica DFC420C camera. All measurements based on photos were performed in ImageJ and analyzed and plotted into graphs in GraphPad Prism 5.

### Phenotypic analysis of reproductive organs

To examine pollen vitality, anthers were collected from flowers just before opening into 70μL Alexander staining buffer [10% ethanol, 0.01% (w/v) Malachite green, 25% glycerol, 5% (w/v) phenol, 5% (w/v) chloral hydrate, 0.05% (w/v) fuchsin acid, 0.005% (w/v) OrangeG and 1.5% glacial acetic acid] on a microscopy slide, covered with cover slip, and incubated at 55 ℃ for 1 hr before imaging. For pollen tube growth, pollen of just opened flowers were transferred to a dialysis membrane placed on solid pollen germination medium [18% Sucrose, 0.01% Boric acid, 1 mM CaCl_2_, 1 mM Ca (NO_3_)_2_, 1 mM MgSO_4_ and 0.5% agarose] and incubated at 22 ℃ for 18 hrs. Ovules were cleared in chloral hydrate solution (chloral hydrate: glycerol: water = 4:2:1 by weight) for 4 hrs. Stained or germinated pollen and cleared ovules were imaged using a Zeiss Axioplan 2 microscope with DIC optics and Zeiss AxioCam MRc 5 digital color camera. Pollen tube length was measured with ImageJ (Fiji).

### Protoplast isolation and transformation

Protoplasts were isolated and transformed as previously described, but with some modifications to the protocol (Schirawski et al., 2000). Protoplasts were isolated from 4-week-old rosette leaves instead of from cell suspensions, and we used a 40% PEG4000 solution and 15 μg *pART7-35S::PID:YFP* for each transformation.

### Auxin transport measurements

Auxin transport assays were carried out as previously reported, with some modifications (Zourelidou et al., 2009). Four 2.5 cm inflorescence stem segments from the basal part of 15cm inflorescence stems were placed in inverted orientation into 30 μL auxin transport buffer (0.5 nM IAA, 1% sucrose, 5 mM MES, pH 5.5) with or without 50 μM NPA for 1 hour, then transferred to 30 μL auxin transport buffer with or without 50 μM NPA containing 200 nM radiolabeled [^3^H]IAA (Scopus Research BV, Veenendaal, The Netherlands), allowed to incubate for 30 minutes and subsequently transferred to 30 μL auxin transport buffer without radiolabeled [^3^H]IAA and incubated for another 4 hrs. Segments were cut into 5 mm pieces, the bottom piece (0-0.5 cm) was discarded and the remaining pieces were placed separately into 5 mL Ultima Gold™ (PerkinElmer, # 6013329) for overnight maceration. The [^3^H]IAA was quantified using a PerkinElmer Tri-Carb 2810TR low activity liquid scintillation analyzer.

### GUS staining and microscopy

Fresh seedlings and plant organs were directly soaked into GUS staining buffer [10 mM EDTA, 50 mM sodium phosphate (pH 7.0), 0.1% (v/v) Triton X-100, 0.5 mM K_3_Fe(CN)_6_, 0.5 mM K_4_Fe(CN)_6_, 1 mg/ml 5-bromo-4-chloro-3-indolyl-D-glucuronide] under vacuum for 15 min and incubated at 37 ℃ for 18 hrs. Subsequently, samples were cleared in 70% (v/v) ethanol at room temperature before imaging with Leica MZ16FA or Leica MZ12 equipped with Leica DFC420C or DC500 camera respectively.

To visualize YFP:PDK1 in embryos and roots or PID:YFP in protoplasts, a Zeiss LSM5 Exciter/AxioImager equipped with a 514 nm laser and a 530-560 nm band pass filter was used on. GFP signals in roots of 5-day-old seedlings were visualized by optionally staining with 10 μg/mL propidium iodide (PI) for 5 min on slides, and observing the samples with a Zeiss LSM5 Exciter/AxioImager equipped with a 488 nm laser and a 505–530 nm band pass filter to detect GFP fluorescence, or a 650 nm long pass filter to detect PI fluorescence. All images were captured with a 40× oil immersion objective (NA = 1.2). Images were optimized in Adobe Photoshop cc2018 and assembled into figures using Adobe Illustrator cc2017. DR5::GFP total intensity was measured from three-dimensional reconstruction of the root tips with ImageJ (Fiji). Apical PIN2:GFP abundance was also measured with ImageJ (Fiji) by drawing a free-hand line along the center of the apical PM of epidermal cells.

### *in vitro* phosphorylation and yeast complementation

*In vitro* phosphorylation and yeast complementation experiments were performed as previously described (Huang et al., 2010; Dittrich and Devarenne, 2012a)

## Acknowledgments

We thank Christian Hardtke and Claus Schwechheimer for providing *pax paxl* and *d6pk012* mutant seeds, respectively. We thank Timothy Devarenne for providing yeast strain *p416GPD*, Sylvia de Pater for providing the CRISPR/Cas9 plasmids, Xiao Men for sharing preliminary data on PIN2HL phosphorylation by PDK1, Kees Boot for help with the auxin transport assay, and Gerda Lamers and Joost Willemse for help with microscopy. We are grateful to Nick Surtel, Ward de Winter and Jan Vink for their help with plant growth and media preparation. This project was supported by the China Scholarship Council.

---

**Figure S1.**
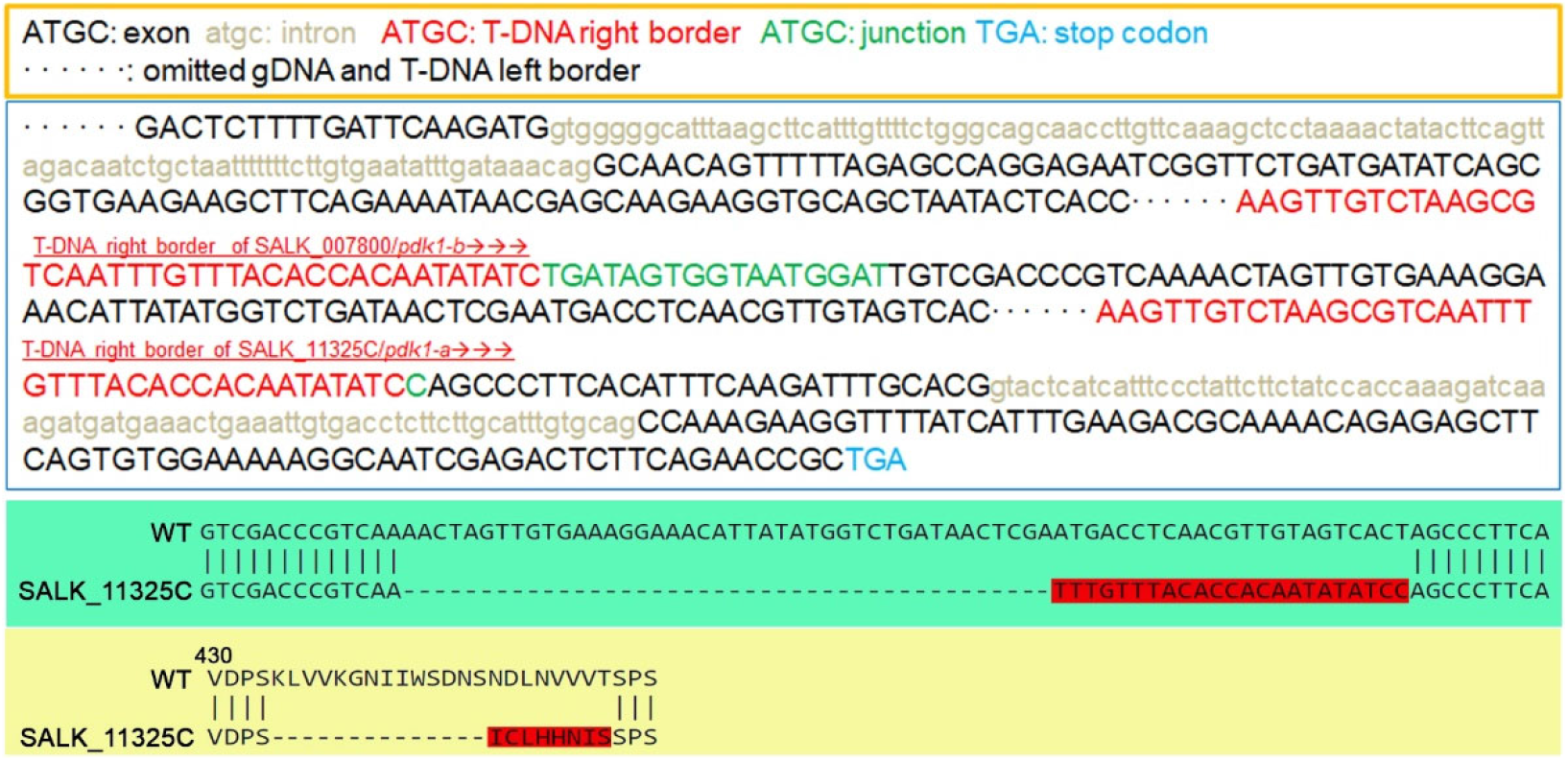
DNA sequence of T-DNA insertion loci in the Arabidopsis *pdk1-a* (SALK_11325) and *pdk1-b* (SALK_007800) T-DNA insertion mutant alleles from NASC. Letters with different colors represent indicated DNA features as indicated in the orange box on the top. Two colored boxes at the bottom show the alignments of part of the PDK1 transcript (green) and the corresponding protein sequence (yellow) in wild-type Arabidopsis (WT) and the *pdk1-a* allele (SALK_11325C). The differences are highlighted in red.

**Figure S2.**
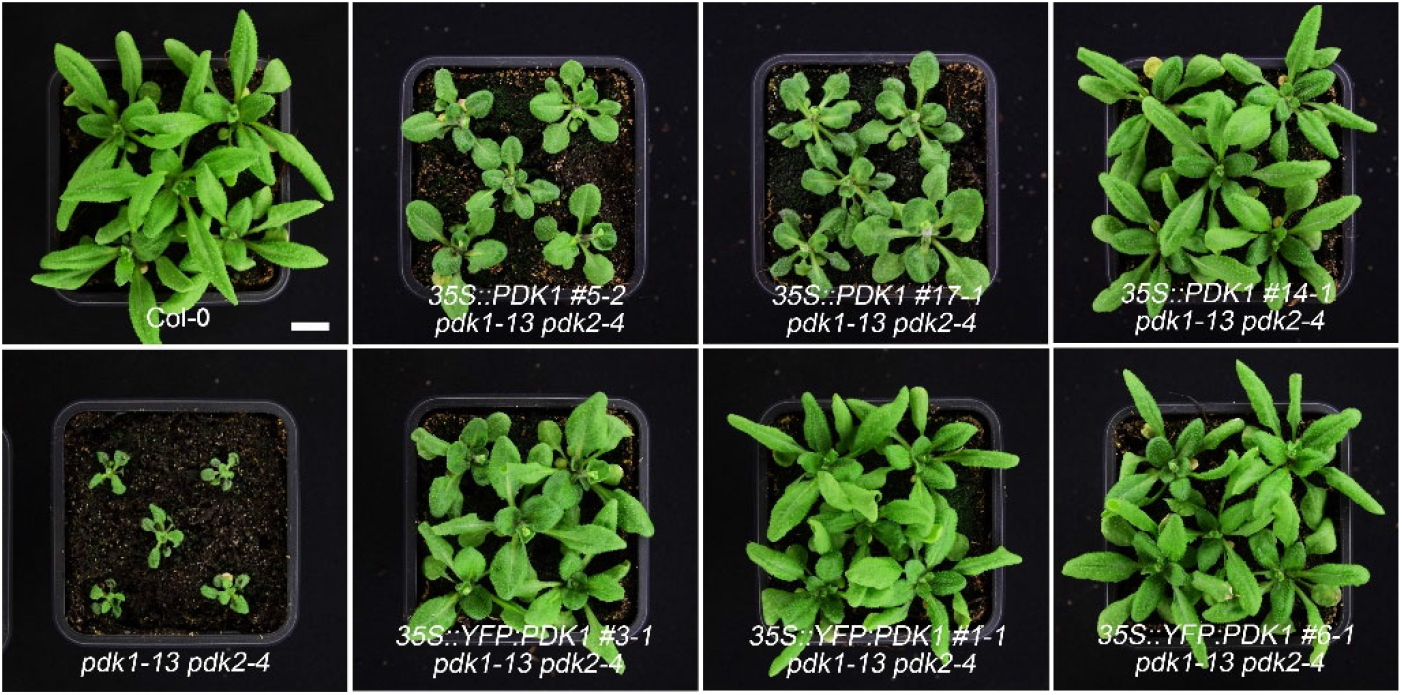
Complementation of *pdk1-13 pdk2-4* by *35S::PDK1* or *35S::YFP:PDK1*. Plants of wild type (Col-0), *pdk1-13 pdk2-4* and representative complementation lines were grown on plates for 10 days then in soil for 20 days before photographing.

**Figure S3.**
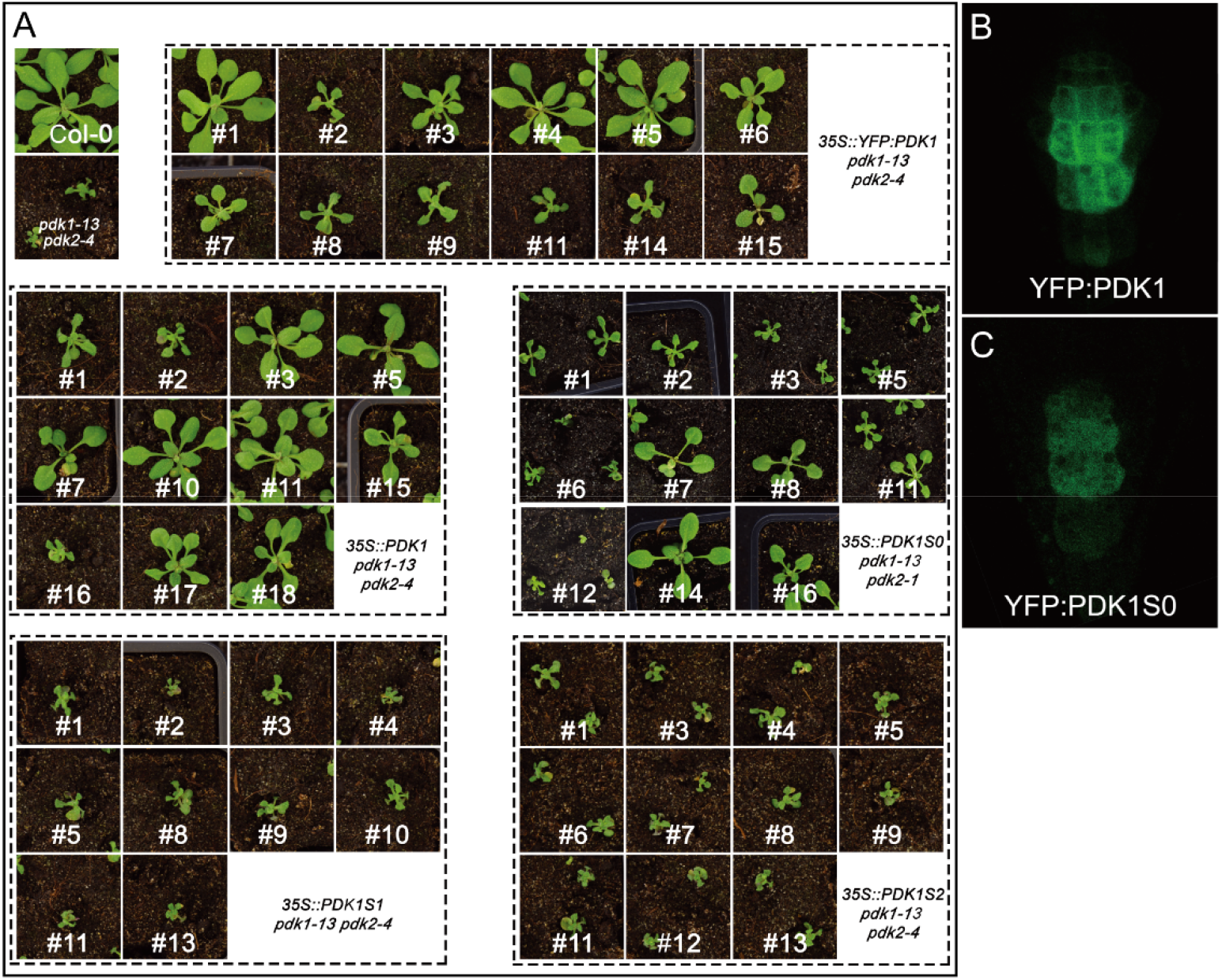
A, Complementation assay for overexpression of cDNAs representing the different *PDK1* splice variants in the *pdk1 pdk2* mutant background. Plants were grown on plates for 10 days, and subsequently transferred to and grown in soil for 10 days. B and C, Subcellular localization of YFP:PDK1 (B) and YFP:PDK1S0 (C) in root columella cells. YFP:PDK1S0 did not show any plasma membrane localization.

**Figure S4.**
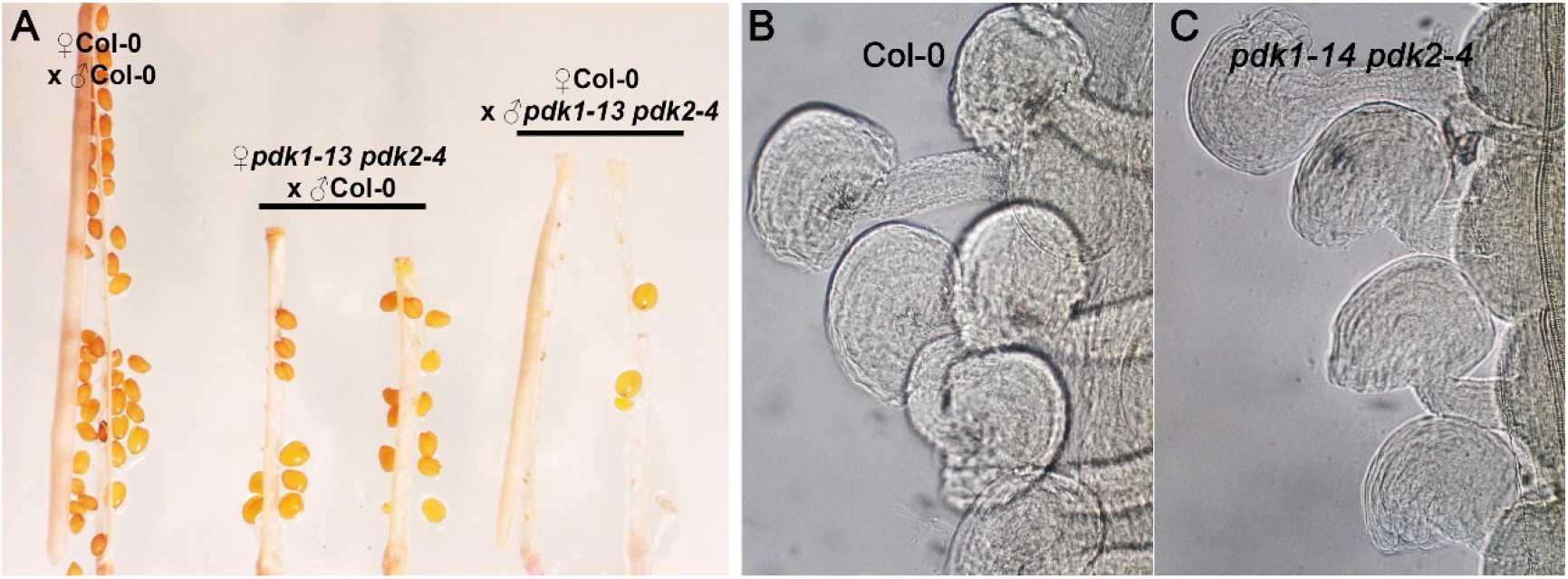
p*d*k1 *pdk2* mutants are strongly defective in male gametophyte development and show normal ovule development. A, Ripe siliques with the valves removed, derived from reciprocal crosses between wild-type Arabidopsis (Col-0) and the *pdk1-13 pdk2-4* loss-of-function mutant B, C, Representative DIC images showing the phenotype of wild-type (B, Col-0, (n>300) and *pdk1-14 pdk2-4* mutant (C, n>300) ovules.

**Figure S5.**
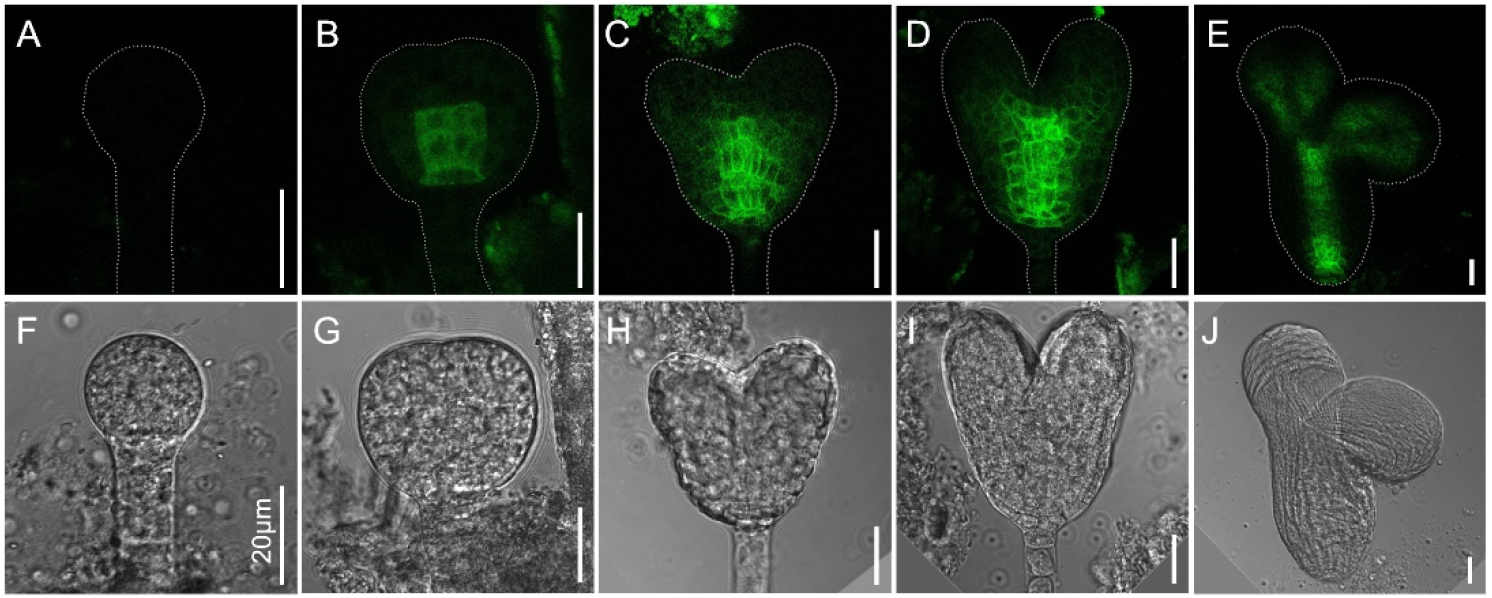
P*D*K1 expression is confined to the provascular tissue during Arabidopsis embryo development. A-E, Confocal (A-E), and bright field images (F-J) of Arabidopsis *pPDK1::YFP:PDK1* 16-cell (A, F), globular (B,G), heart (C,H), late heart (D,I), and torpedo (E,J) stage embryos.

**Figure S6.**
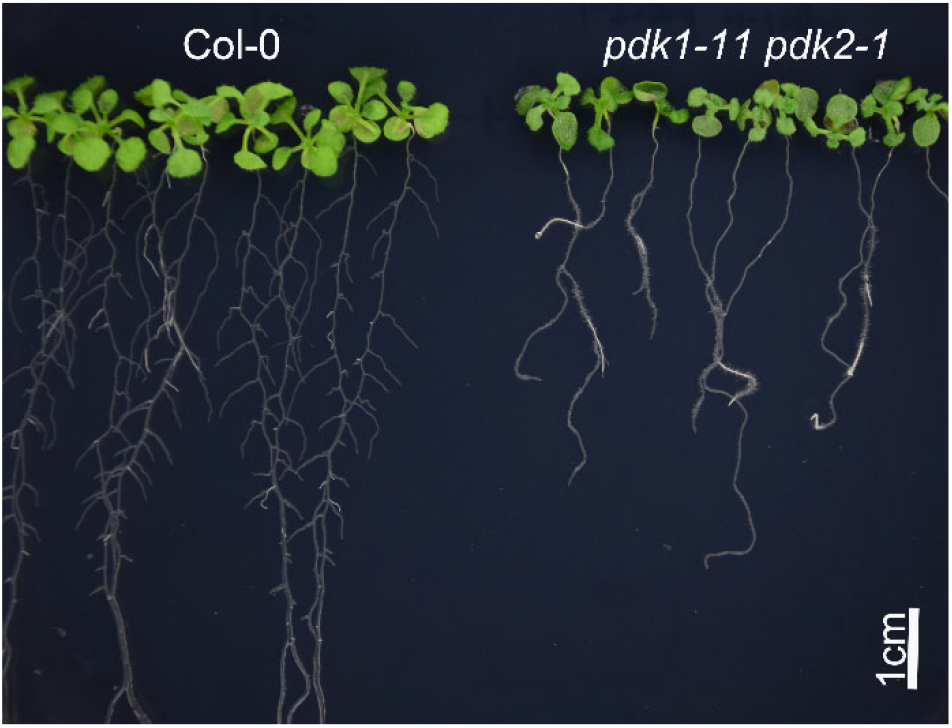
Phenotype of 15-day-old wild-type (Col-0) and *pdk1-11 pdk2-1* seedlings grown on vertical plates.

**Table S1.** Frequency of *pdk1 pdk2* double homozygous progeny obtained from *pdk1* (-/-) *pdk2* (+/-) (green) or *pdk1* (+/-) *pdk2* (-/-) (yellow) parent plants. n>130. n. a.: not analysed.

|  | *pdk2-1* | *pdk2-4* |
| --- | --- | --- |
| *pdk1-11* | 1:5.8 | 1:9.6 |
| *pdk1-13* | 1:7.3 | 1:13.3 |
| *pdk1-14* | 1:26.6 | 1:12.3 |
| *pdk1-31* | n.a. | 1:9.3 |
| *pdk1-32* | 1:7.5 | 1:5.7 |

